# Integrated Transcriptomic and Translatomic Profiling Reveals Noncanonical RNA Junctions and Hidden Coding Potential in Porcine Epidemic Diarrhea Virus

**DOI:** 10.64898/2026.11.05.764755

**Authors:** Guiwen Guan, Ran Zhang, Jin Wang, Hanwen Zhang, Leyu Hu, Peixuan Li, Xuanhao Xu, Mingjun Zhu, Qian Du, Lu Han, Xin Ran, Yong Huang

## Abstract

Porcine epidemic diarrhea virus (PEDV) poses a major threat to the global swine industry. Although coronavirus discontinuous transcription is well characterized, the classical model cannot fully explain the diversity of viral RNA junctions, while conventional genome annotations may underestimate viral protein-coding capacity. To address these gaps, we integrated time-resolved RNA sequencing (RNA-seq), cycloheximide (CHX)- and harringtonine (HAR)-treated ribosome profiling (Ribo-seq), comparative genomics, and peptide mass spectrometry to investigate PEDV RNA expression and hidden coding potential.

RNA-seq revealed that, at 3 hours post-infection (hpi), the 3’ genomic region exhibited strong relative coverage enrichment, whereas the relative contribution of the 5’ replicase-coding region increased rapidly between 3 and 5 hpi, suggesting a possible transition from sgRNA-dominated RNA accumulation toward enhanced genomic RNA replication. Genome-wide junction analysis identified 43,635 distinct noncanonical RNA junctions, although canonical sgRNA junctions accounted for 95.5% of all junction-supporting reads. Several abundant noncanonical sgRNAs were predicted to generate novel coding products, including N-x4, N-x2, M-x3, and M-x2, short peptides encoded within the 3’ untranslated region (UTR), and two distinct N-terminally truncated S proteins, thereby expanding the potential coding repertoire of PEDV.

Genome-wide reading-frame analysis uncovered a pronounced asymmetry in hidden coding potential, identifying 68 overlapping open reading frames (ORFs) longer than 40 amino acids in the +1 reading frame, compared with only three in the +2 frame. Ribo-seq further revealed a progressive redistribution of viral ribosome occupancy toward structural protein-coding regions during the later stages of infection. By integrating HAR-associated translation initiation-site enrichment with CHX-based ORF-level translation analysis, we identified several noncanonical ORFs with strong translation-associated evidence, including N-x1, N-x2, S-x1, and ORF1b-x13. Moreover, peptide mass spectrometry detected a characteristic peptide corresponding to N-x1 in infected samples, providing independent proteomic evidence for the expression of this previously unannotated viral protein.

Together, these findings uncover extensive noncanonical RNA synthesis and hidden protein-coding capacity in PEDV, establishing a foundation for understanding novel viral gene expression strategies and their potential evolutionary and pathogenic significance.

## Introduction

Porcine epidemic diarrhea virus (PEDV) is a major causative agent of viral enteric disease in swine and continues to pose a substantial threat to the global pig industry. Despite widespread vaccination and biosecurity measures, PEDV remains prevalent in multiple regions ^1,2^. Different PEDV strains exhibit considerable variation in replication capacity, tissue tropism, and pathogenicity ^3,4^, which cannot be fully explained by the known functions and sequence variations of annotated viral proteins. Viral replication and pathogenesis depend not only on the functions of encoded proteins but also on the precise regulation of viral RNA synthesis and protein expression^1,3,4^. Therefore, a deeper understanding of PEDV gene expression strategies at the transcriptional and translational levels is essential for elucidating its replication and pathogenic mechanisms and identifying potential avenues for intervention.

As an alphacoronavirus, PEDV possesses a positive-sense, single-stranded RNA genome of approximately 28 kb^1^. Its replication and gene expression involve genome replication, direct translation of genomic RNA, and the production of subgenomic RNAs (sgRNAs) through discontinuous transcription^5^. The formation of sgRNAs involves template switching by the viral RNA-dependent RNA polymerase during negative-strand synthesis and is influenced by sequence complementarity between the leader transcription-regulating sequence (TRS-L) and downstream body transcription-regulating sequences (TRS-B), as well as local RNA structures^5–7^. Recent high-resolution transcriptomic studies have demonstrated that the repertoire of coronavirus RNAs is substantially more complex than that described by the classical model of nested sgRNAs^8,9^. In addition to canonical sgRNAs, coronaviruses can generate noncanonical leader–body junctions, internal genomic junctions, and other discontinuous RNA species^8,9^. In PEDV, the relative abundance, exact breakpoint heterogeneity, and infection-associated dynamics of these RNA junctions remain incompletely characterized. More importantly, the biological implications of noncanonical RNA junctions may extend beyond transcript architecture. Discontinuous joining of different genomic regions may truncate canonical coding sequences, alter reading-frame relationships across junctions, or generate new open reading frames (ORFs), thereby expanding the potential coding capacity of viral RNAs^8,9^. Thus, systematic characterization of canonical and noncanonical PEDV RNA junctions may not only improve our understanding of viral RNA synthesis and its sequence-dependent regulation but also reveal changes in coding potential associated with noncanonical RNA architectures, providing a basis for exploring previously unrecognized viral expression products.

However, viral RNA abundance and predicted coding potential do not necessarily reflect actual protein synthesis. Ribosome profiling (Ribo-seq), which captures ribosome-protected fragments, enables the analysis of translation activity at near-codon resolution and provides direct evidence of ribosome occupancy, reading-frame usage, and infection-stage-dependent changes in viral protein expression^10,11^. Recent Ribo-seq studies of other coronaviruses have revealed dynamic translation patterns across viral coding regions and identified multiple translation-associated regions beyond conventional genome annotations, including upstream short ORFs, alternative translation initiation sites, and ORFs overlapping canonical coding sequences^11,12^. These findings suggest that coronaviruses may further expand their coding capacity through noncanonical translation strategies^11^. Although the major PEDV proteins and certain aspects of their translational regulation have been investigated, a systematic understanding of translation dynamics across canonical coding regions, genome-wide reading-frame usage, and the actual translation of predicted noncanonical ORFs during PEDV infection remains lacking. In particular, numerous potential alternative ORFs in the PEDV genome overlap canonical coding sequences, where strong canonical translation signals may obscure low-abundance ribosome occupancy in alternative reading frames. Consequently, sequence-based predictions and localized ribosome enrichment alone may be insufficient to establish reliable evidence of noncanonical translation. Integrating translation initiation-associated ribosome enrichment, frame-resolved translation analysis, and independent proteomic evidence is therefore particularly important for identifying PEDV noncanonical ORFs with genuine expression potential^11,13^. Such an approach may not only refine the PEDV protein-coding landscape but also provide new candidates for investigating viral factors potentially involved in replication, host adaptation, and pathogenesis.

To address these questions, we integrated time-resolved RNA sequencing (RNA-seq), ribosome profiling following cycloheximide (CHX) or harringtonine (HAR) treatment, comparative analysis of PEDV genomes, and peptide mass spectrometry to systematically characterize viral RNA architectures and translation-associated activities during PEDV infection. By examining RNA structures, ribosome occupancy, and proteomic evidence, this study aims to expand our understanding of PEDV gene expression strategies and provide a molecular foundation for investigating transcriptional and translational regulation during coronavirus replication, as well as the potential biological roles of previously unannotated viral coding products.

### Results 1. PEDV RNA accumulation exhibits asynchronous regional dynamics and MOI-dependent changes during infection

To characterize the accumulation and genomic distribution of PEDV RNA during infection, Vero cells were infected with a PEDV GI-b strain at multiplicities of infection (MOIs) of 0.1 or 1 and collected at 3, 5, 8, 12, and 16 hours post-infection (hpi) for RNA sequencing (RNA-seq) (Fig 1A). Western blot analysis of viral nucleocapsid (N) protein and RT-qPCR measurements confirmed progressive viral accumulation during infection, while sequencing quality assessment, sample correlation analysis, and principal component analysis supported the reliability of the datasets (Fig S1 and S2). In addition, to validate our findings in porcine tissues, porcine-derived cells, and across different PEDV genotypes, we retrieved publicly available RNA-seq datasets from the Sequence Read Archive (SRA), including datasets generated from jejunum of PEDV-infected piglets and LLC-PK1 cells infected with a PEDV GII-b strain.

**Fig 1.**
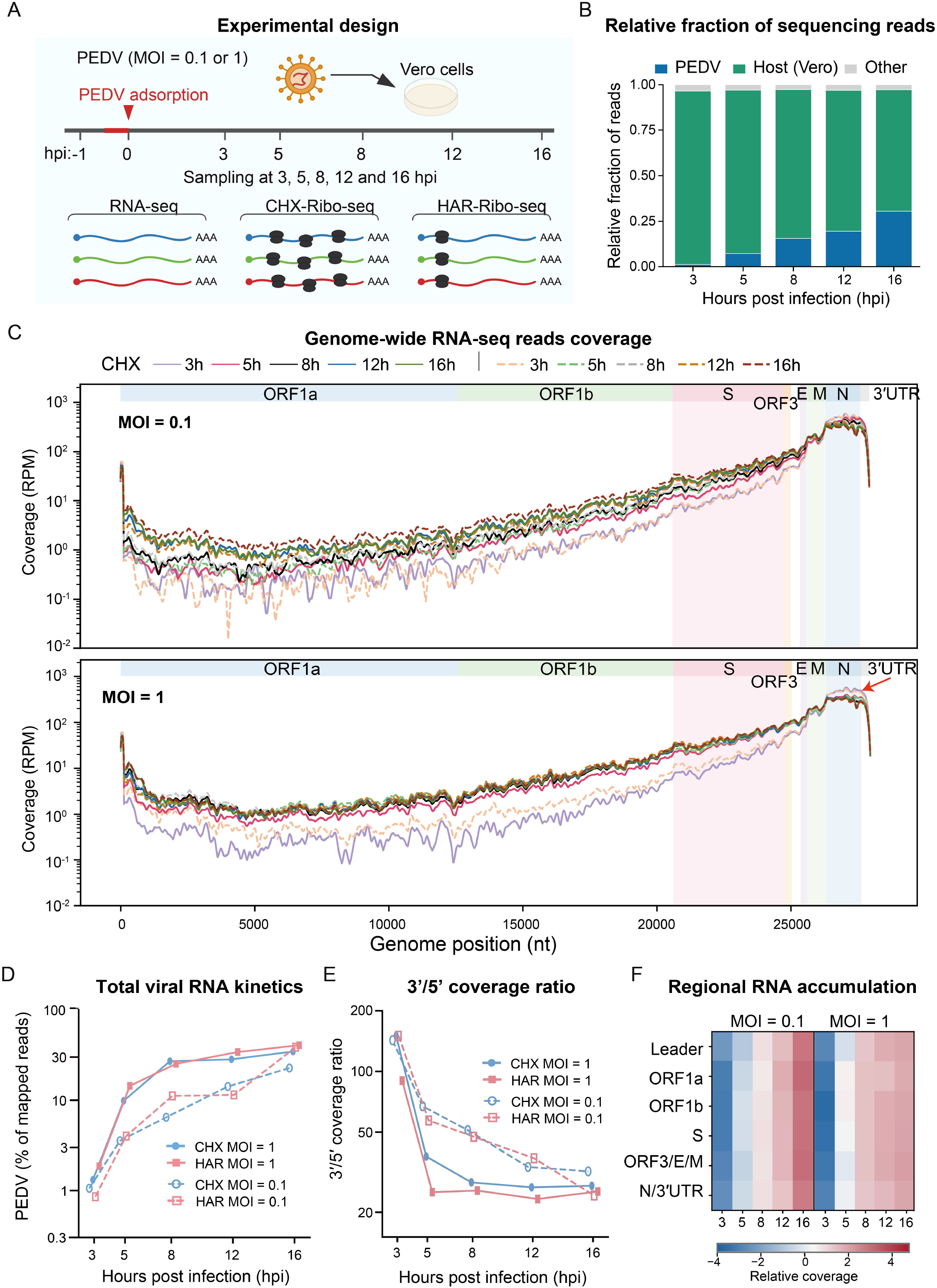
Time-resolved RNA-seq reveals dynamic and regionally heterogeneous PEDV RNA accumulation during infection. (A) Experimental design. Vero cells were infected with PEDV at multiplicities of infection (MOIs) of 0.1 or 1 and sampled at 3, 5, 8, 12, and 16 hours post-infection (hpi). RNA-seq and Ribo-seq libraries were generated from cycloheximide (CHX)- and harringtonine (HAR)-treated samples. (B) Relative proportions of sequencing reads assigned to PEDV, host, and other sources across infection time points. (C) Genome-wide PEDV RNA-seq coverage at MOI 0.1 and MOI 1. Coverage is expressed as reads per million (RPM) and plotted on a logarithmic scale. Viral genomic regions are indicated above each panel. (D) Temporal changes in the proportion of sequencing reads mapped to PEDV under different MOI and treatment conditions. (E) Ratio of RNA-seq coverage between the genomic 3’ and 5’ regions across infection time points. (F) Heatmap showing regional PEDV RNA accumulation across MOIs and infection times. Values represent row-wise z-scores of normalized coverage for each genomic region.

As infection progressed, the proportion of sequencing reads derived from PEDV increased continuously, accompanied by a corresponding decrease in host-derived reads (Fig 1B). Viral RNA accumulation exhibited pronounced MOI-dependent kinetics. At an MOI of 1, the proportion of viral reads increased rapidly between 3 and 8 hpi, followed by a slower rate of increase. In contrast, at an MOI of 0.1, viral RNA accumulated more gradually during early infection and continued to increase through 16 hpi (Fig 1D and S2E). Despite the overall increase in viral RNA abundance, sequencing coverage varied substantially across genomic regions. At both MOIs, coverage was relatively low across ORF1a and ORF1b and progressively increased toward the genomic 3’ end, reaching its highest levels in the N gene and adjacent regions. Coverage in the N region was approximately 100- to 1,000-fold higher than that in ORF1a (Fig 1C). This distribution is consistent with the accumulation of shared 3’-terminal sequences characteristic of nested coronavirus subgenomic RNAs (sgRNAs), although the extent of 3’ enrichment changed markedly over the course of infection^5^.

Notably, despite the relatively low overall abundance of viral RNA at 3 hpi, the ratio of 3’-to-5’ genomic coverage reached its maximum of approximately 150 and subsequently declined rapidly as viral RNA accumulated (Fig 1E). At an MOI of 1, this ratio decreased to approximately 25–35 by 5 hpi and remained relatively low thereafter. In contrast, the decline was slower at an MOI of 0.1, indicating that pronounced relative enrichment of 3’-terminal sequences persisted for a longer period under the lower inoculation condition. Importantly, this change does not indicate a reduction in the absolute abundance of 3’-terminal viral RNA. Rather, it suggests that sgRNA-associated sequences may make a greater relative contribution during early infection, whereas the subsequent increase in replicase-associated genomic sequences may reflect accelerated genomic RNA replication after 3 hpi. Thus, the declining 3’/5’ coverage ratio is consistent with a temporal shift in the relative contributions of viral transcription and genome replication. Regional analysis of the leader, ORF1a, ORF1b, S, ORF3/E/M, and N/3’UTR regions further supported these dynamic changes (Fig 1F).

Together, these results reveal temporal changes in the relative contributions of genomic RNA-associated and structural protein sgRNA-associated sequences during PEDV infection, providing a basis for subsequent junction-resolved characterization of canonical and noncanonical viral RNA species.

### Results 2. PEDV RNA junctions reveal extensive noncanonical transcription and position-dependent template-switching patterns

To further characterize the composition of viral RNAs during PEDV infection and explore the mechanisms underlying their formation, we systematically identified RNA-seq reads spanning discontinuous RNA junctions and mapped their corresponding 5’ and 3’ breakpoints to the viral genome (Fig 2A and S3). Genome-wide two-dimensional breakpoint mapping revealed that PEDV RNA junctions were not restricted to a small number of canonical sgRNA sites but were widely distributed across the viral genome, exhibiting pronounced spatial heterogeneity (Fig 2A). High-abundance junctions involving the 5’ leader region predominantly corresponded to canonical sgRNAs encoding structural and accessory proteins, whereas numerous low-abundance junctions connected other genomic regions, generating diverse long-range and local RNA junctions.

**Fig 2.**
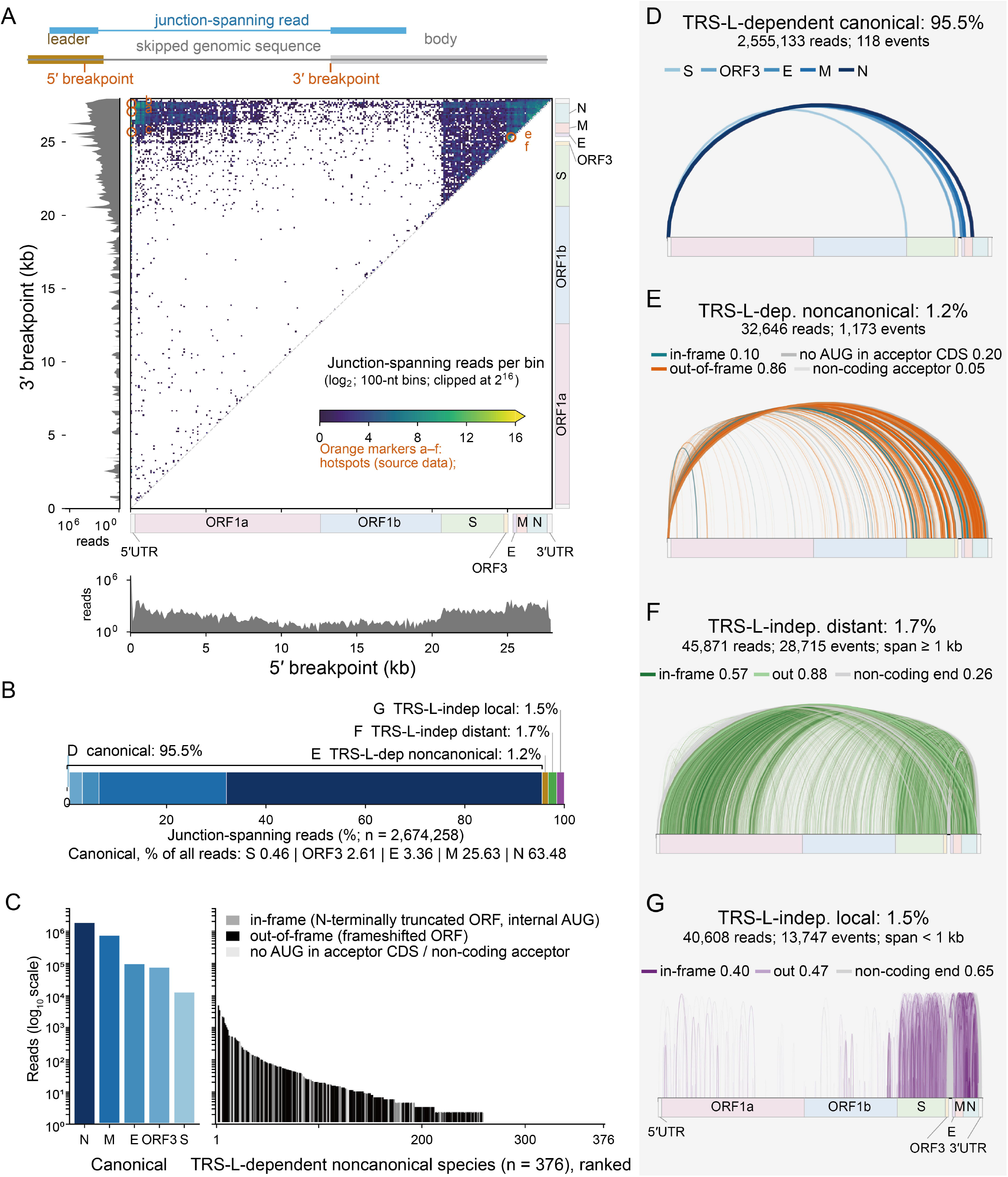
Genome-wide mapping reveals extensive noncanonical RNA junction diversity dominated by canonical PEDV sgRNAs. **(A)** Genome-wide two-dimensional distribution of RNA junction breakpoints. The x- and y-axes represent the 5’ and 3’ breakpoints, respectively. Junction-supporting reads were aggregated into 100-nt bins and displayed on a log□ scale, with marginal distributions showing read counts at each breakpoint position. Orange circles indicate representative junction hotspots. **(B)** Relative abundance of four junction classes among 2,674,258 classified junction-supporting reads. **(C)** Read abundance of the five canonical sgRNAs and 376 TRS-L-dependent noncanonical RNA species, ranked by abundance. **(D)** Genomic connections corresponding to canonical leader–body junctions, accounting for 95.5% of classified junction-supporting reads. **(E)** TRS-L-dependent noncanonical junctions (1.2%), colored according to their predicted coding consequences. **(F)** TRS-L-independent distant junctions (≥1 kb; 1.7%), categorized by their potential coding consequences. **(G)** TRS-L-independent local junctions (<1 kb; 1.5%), categorized by their potential coding consequences. Percentages in panels D–G indicate the proportion of all classified junction-supporting reads.

Examination of the marginal breakpoint distributions revealed distinct spatial patterns. The 3’ breakpoints were strongly enriched toward the genomic 3’ end, with the number of supporting reads generally decreasing in the 3’-to-5’ direction and prominent local peaks occurring near canonical body transcription-regulating sequences (TRS-Bs) (Fig 2A and S3). In contrast, 5’ breakpoints were highly concentrated in the leader region but also showed enrichment near the genomic 3’ end and at several internal positions, whereas junction density was relatively low across the central ORF1a/ORF1b regions (Fig 2A and S3). These breakpoint distributions prompted us to propose a possible mechanistic explanation. During negative-strand RNA synthesis, as the PEDV polymerase progresses along the positive-strand template from its 3’ end toward its 5’ end, low-probability template dissociation and rejoining events may occur continuously in addition to canonical TRS-mediated template switching^5,6^. As synthesis advances toward the genomic 5’ end, the number of polymerases remaining engaged in elongation on the original template may progressively decline, potentially producing the overall decreasing background gradient of 3’ breakpoint-supporting reads. Near canonical TRS-B sites, specific sequence features and local RNA structures may facilitate polymerase pausing or template switching, giving rise to prominent junction hotspots above this background. Meanwhile, strong sequence complementarity between the nascent negative-strand RNA and the leader transcription-regulating sequence (TRS-L) may favor rejoining at the leader region, whereas some intermediates that do not rejoin the leader may instead reanneal to other genomic positions, generating noncanonical distant or local junctions^5,6^.

Despite the extensive diversity of RNA junction architectures, canonical sgRNAs overwhelmingly dominated junction-supporting read abundance. Among 2,674,258 classified junction-spanning reads, 2,555,133 (95.5%) corresponded to canonical TRS-L-dependent junctions (Fig 2B–D). A similar proportion of canonical junctions was also observed in RNA-seq datasets from the jejunum of PEDV GII-b-infected piglets and PEDV GII-b-infected LLC-PK1 cells (S4A and S4B). Notably, the predominance of canonical sgRNAs did not imply that their junction boundaries were completely invariant. Analysis of exact breakpoint coordinates identified 10–47 distinct junction variants within each of the five canonical sgRNA classes (Fig S5A) For S, ORF3, E, and N, the most abundant exact junction accounted for 97.6%, 97.7%, 97.2%, and 99.6% of the corresponding class-specific reads, respectively, indicating highly focused breakpoint selection. In contrast, M sgRNA exhibited a distinctly heterogeneous junction pattern, with three major junctions, 66→ 25597, 63→25594, and 67→25598, accounting for 42.4%, 31.5%, and 25.8% of M-associated junction reads, respectively (Fig S5B).

To systematically characterize the diversity of discontinuous RNA connections, we classified junctions into four categories according to their association with TRS-L and genomic span: TRS-L-dependent canonical junctions, TRS-L-dependent noncanonical junctions, TRS-L-independent distant junctions (span ≥1 kb), and TRS-L-independent local junctions (span <1 kb) (Fig 2D-G). In addition to 118 canonical exact junction events, we identified 43,635 noncanonical events, comprising 1,173 TRS-L-dependent noncanonical junctions, 28,715 distant junctions, and 13,747 local junctions (Sup table 1). These three noncanonical classes accounted for only 1.2%, 1.7%, and 1.5% of all classified junction-supporting reads, respectively. Thus, although noncanonical junctions represented more than 99% of distinct exact junction events, they contributed only approximately 4.5% of junction-supporting reads. This striking contrast highlights an important feature of PEDV RNA populations: junction architectures are highly diverse, but the abundance of detectable junction products is strongly concentrated at a limited number of canonical sites.

We next examined how infection time and MOI influenced junction composition. Canonical junctions remained dominant across all experimental conditions (Fig S6A). In contrast, the relative composition of noncanonical junctions exhibited more pronounced temporal variation. TRS-L-dependent noncanonical junctions remained relatively stable, whereas TRS-L-independent distant and local junctions fluctuated across infection stages (Fig S6B). These changes were particularly evident at an MOI of 1, where the relative contributions of some noncanonical classes were higher during early infection and subsequently declined or partially recovered (Fig S7). Similar class-specific temporal differences were observed after normalization to the total number of viral fragments (Fig S6C).

We further examined the correlations between noncanonical sgRNA junction profiles identified in Vero cells infected with PEDV GI-b and those detected in jejunum from PEDV-infected piglets and LLC-PK1 cells infected with PEDV GII-b. The junction profiles showed a relatively strong correlation between Vero and LLC-PK1 cells (Spearman ρ = 0.60, *P* < 0.001), suggesting a degree of conservation in PEDV RNA junction patterns across different cell types and viral genotypes. In contrast, the correlation between Vero cells and porcine jejunum was relatively weak (Spearman ρ = 0.24). Further analysis indicated that this limited correlation was primarily attributable to insufficient PEDV sequencing coverage in the jejunal RNA-seq dataset (Fig S4C and S4D).

Finally, we evaluated the potential coding consequences of TRS-L-dependent noncanonical sgRNAs. Based on their predicted ORFs, these transcripts were categorized into three groups: sgRNAs potentially encoding N-terminally truncated canonical proteins, sgRNAs potentially encoding alternative-frame ORFs, and sgRNAs potentially encoding ORFs within untranslated regions. We focused on the 20 most abundant and recurrent noncanonical sgRNAs (Fig S8A). The most abundant junction, 63→27061, was predicted to support expression of N-x4, a 49-amino-acid alternative-frame ORF within the N coding region. The second most abundant junction, 61→27608, was predicted to support a 35-amino-acid ORF in the 3’ UTR (Fig S8B). Other frequently detected junctions were associated with potential alternative-frame products, including M-x2, M-x3, and N-x2. Notably, junctions 56→22456 and 60→21491 were predicted to generate N-terminally truncated S proteins of 1,084 and 743 amino acids, respectively. All of these highly abundant noncanonical sgRNA junctions were also detected in PEDV GII-b-infected LLC-PK1 cells. In contrast, only a subset of these junctions was detected in the jejunal tissue RNA-seq dataset from PEDV-infected piglets. We speculate that this limited detection may also be largely attributable to insufficient PEDV sequencing coverage in the porcine jejunal tissue dataset (Fig S9A-C). These findings indicate that noncanonical leader–body junctions may expand the coding potential of PEDV RNAs by generating alternative transcript architectures, although translation of the corresponding predicted products remains to be experimentally established.

### Results 3. PEDV RNA junctions exhibit distinct degrees of TRS-L sequence similarity and local microhomology

To investigate the sequence features potentially underlying the formation of different classes of PEDV RNA junctions, we first compared the local sequence characteristics of the leader transcription-regulating sequence (TRS-L) with those of the five canonical body transcription-regulating sequences (TRS-Bs) associated with S, ORF3, E, M, and N. The conserved PEDV TRS core was identified as CTAAAC, which was fully retained in the TRS-B of the N sgRNA, whereas the TRS-B core sequences of the remaining canonical sgRNAs exhibited varying degrees of mismatch with TRS-L. This contrasts with the more uniform TRS-B core sequences commonly observed in some other coronaviruses^5,6^. Nevertheless, all five canonical TRS-Bs displayed recognizable sequence similarity to TRS-L, although the extent and positions of matching nucleotides varied among them (Fig 3A). Consistent with these features, canonical junction acceptor sites exhibited concentrated nucleotide preferences and pronounced peaks of TRS-L similarity (Fig 3B and 3C), corresponding to the strong enrichment of canonical sgRNAs at a limited number of preferred junction sites. Notably, although the 4,073 noncanonical junction acceptor sites lacked the strong and sharply localized sequence conservation observed at canonical TRS-Bs, they retained a detectable degree of TRS-L-related sequence preference that differed from the background represented by 83,610 random genomic positions (Fig 3B). This weaker local similarity may be particularly relevant to TRS-L-dependent noncanonical sgRNAs. Although most noncanonical sites lack the extensive complementarity characteristic of canonical TRS-Bs, short or imperfect sequence matches at some sites may facilitate reannealing of the nascent negative-strand RNA to the 5’ leader region, thereby allowing the formation of low-abundance noncanonical leader – body junctions. In contrast, TRS-L-independent distant and local junctions do not involve leader joining and therefore cannot be directly explained by TRS-L similarity.

**Fig 3.**
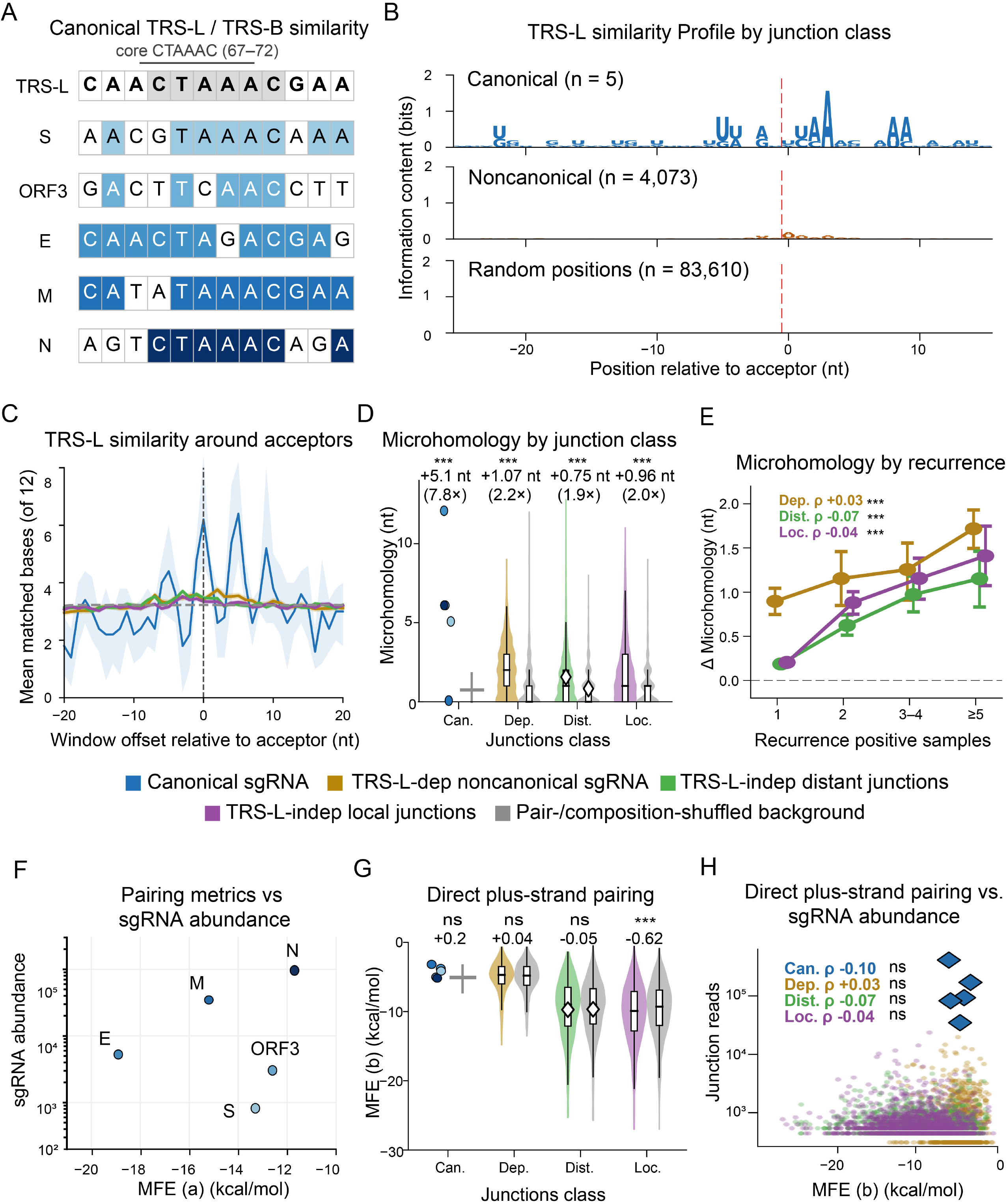
TRS-L sequence similarity and junction microhomology characterize distinct classes of PEDV RNA connections. **(A)** Sequence similarity between the canonical leader transcription-regulating sequence (TRS-L) and body transcription-regulating sequences (TRS-B) associated with the five canonical sgRNAs. Shading indicates matching nucleotides. **(B)** Sequence information profiles surrounding canonical and noncanonical junction acceptor sites compared with random genomic positions. Position 0 indicates the junction acceptor. **(C)** Mean sequence similarity to TRS-L across 12-nt windows positioned relative to junction acceptors. Lines represent different junction classes, with the shaded region indicating variability for canonical sites. **(D)** Junction microhomology lengths compared with pair- and sequence composition-shuffled backgrounds. Labels indicate the mean enrichment relative to the corresponding background. **(E)** Microhomology enrichment relative to shuffled backgrounds across groups defined by the number of samples supporting each noncanonical junction. Points and error bars summarize microhomology enrichment within recurrence groups. **(F)** Relationship between predicted RNA pairing MFE and junction-supporting read abundance for the five canonical sgRNAs. **(G)** Predicted minimum free energy (MFE) of direct plus-strand RNA pairing across junction classes compared with shuffled backgrounds. Labels indicate the difference in MFE between observed and background sequences. **(H)** Relationship between predicted direct plus-strand pairing MFE and junction-supporting read abundance across junction classes. Spearman correlation coefficients are indicated in the panel. Statistical annotations: ns, not significant; ***, *P* < 0.001.

Because noncanonical internal junctions connect two genomic positions outside the leader region, we next examined short-sequence microhomology between junction breakpoints and compared the observed values with matched sequence-shuffled backgrounds (Fig 3D). Canonical sgRNA junctions exhibited significant but heterogeneous microhomology enrichment. Importantly, all three noncanonical junction classes also showed microhomology levels exceeding their corresponding backgrounds. Mean microhomology was increased by 1.07 nucleotides (nt) for TRS-L-dependent noncanonical junctions, 0.75 nt for TRS-L-independent distant junctions, and 0.96 nt for TRS-L-independent local junctions, with statistically significant enrichment in all three classes. These findings indicate that short-sequence matching may represent a shared feature of different junction types, despite differences in their joining positions. We further examined whether microhomology enrichment was associated with the recurrence of junctions across independent samples (Fig 3E). Although some more frequently detected junction groups exhibited greater average microhomology enrichment, correlations between microhomology and recurrence at the individual-junction level were weak. Thus, short-sequence complementarity may contribute to sequence-dependent junction selection but does not, by itself, adequately explain the recurrence of individual junctions.

We subsequently investigated whether predicted RNA pairing stability could further account for junction formation and abundance. The five canonical sgRNAs differed in junction-supporting read abundance by several orders of magnitude, yet their abundances showed no clear monotonic relationship with the predicted minimum free energy (MFE) of RNA pairing (Fig 3F). Compared with sequence composition-matched shuffled backgrounds, canonical junctions, TRS-L-dependent noncanonical junctions, and TRS-L-independent distant junctions showed no significant differences in predicted direct plus-strand RNA pairing energy. Only TRS-L-independent local junctions exhibited a modest but statistically significant reduction in MFE (ΔMFE ≈ −0.62 kcal/mol), suggesting slightly greater predicted pairing stability in their local sequences (Fig 3G). Furthermore, predicted pairing energies were not significantly correlated with junction-supporting read abundance in any junction class (Fig 3H). These results indicate that, under the pairing model used in this study, RNA duplex thermodynamic stability alone is a poor predictor of the relative abundance of junction products.

Together, these findings reveal distinct sequence-selection features among PEDV RNA junction classes. Canonical sgRNAs are associated with relatively strong TRS-L/TRS-B sequence matching, whereas a subset of noncanonical leader–body sgRNAs may exploit weaker local complementarity to facilitate leader joining. In contrast, leader-independent internal junctions generally exhibit breakpoint microhomology exceeding the sequence-shuffled background. These observations support a potential role for short-sequence complementarity in different forms of template rejoining, while indicating that sequence matching alone cannot explain the substantial differences in abundance among RNA junction products.

### Results 4. Ribosome profiling reveals temporal redistribution of PEDV translation-associated signals and reading-frame-specific occupancy

To investigate translational activity and its temporal dynamics during PEDV infection, we performed ribosome profiling (Ribo-seq) at 5, 8, and 16 hours post-infection (hpi) following treatment with cycloheximide (CHX) or harringtonine (HAR), providing complementary information on elongating ribosomes and translation initiation-associated ribosome enrichment^10^. Following adapter and fragment-length filtering, removal of rRNA- and noncoding RNA-derived reads, and alignment against host and viral reference genomes, substantial numbers of PEDV-derived ribosome-protected fragments (RPFs) were retained in each library (Fig S10A). Under CHX treatment, the number of strand-assignable viral reads increased from 1.62 million at 5 hpi to 5.84 million at 16 hpi, whereas HAR-treated libraries showed an increase from 1.11 million to 9.38 million over the same period (Fig S10B). Most PEDV-derived reads mapped to the positive strand, although a smaller proportion of negative-strand-associated protected fragments was also detected, with their relative abundance declining markedly at 8 and 16 hpi (Fig S10B and S10C).

To evaluate the reliability of the viral Ribo-seq datasets, we first examined the characteristic features of RPFs. Compared with RNA-seq, Ribo-seq reads were strongly enriched within annotated coding sequences (CDSs) and substantially depleted in 3’ untranslated regions (UTRs) (Fig 4A). Both host- and PEDV-derived RPFs were predominantly 28–31 nucleotides (nt) long, although their length distributions were not identical, with some PEDV libraries exhibiting a higher proportion of 29–31-nt fragments (Fig 4B, S10D and S10E). Following P-site offset calibration using annotated host CDSs, host RPFs displayed pronounced enrichment in the annotated frame 0, whereas RNA-seq reads exhibited an approximately uniform reading-frame distribution (Fig 4C). Metagene analysis further revealed characteristic P-site enrichment around annotated start codons, with HAR-treated libraries displaying particularly sharp initiation-associated peaks and CHX-treated libraries exhibiting broader distributions consistent with elongating ribosomes. Both treatments also showed the expected three-nucleotide periodicity around annotated stop codons and a reduction in P-site signals downstream of termination sites (Fig 4D). Together, these features support the use of the resulting RPF datasets to investigate host and viral ribosome occupancy at nucleotide and reading-frame resolution.

**Fig 4.**
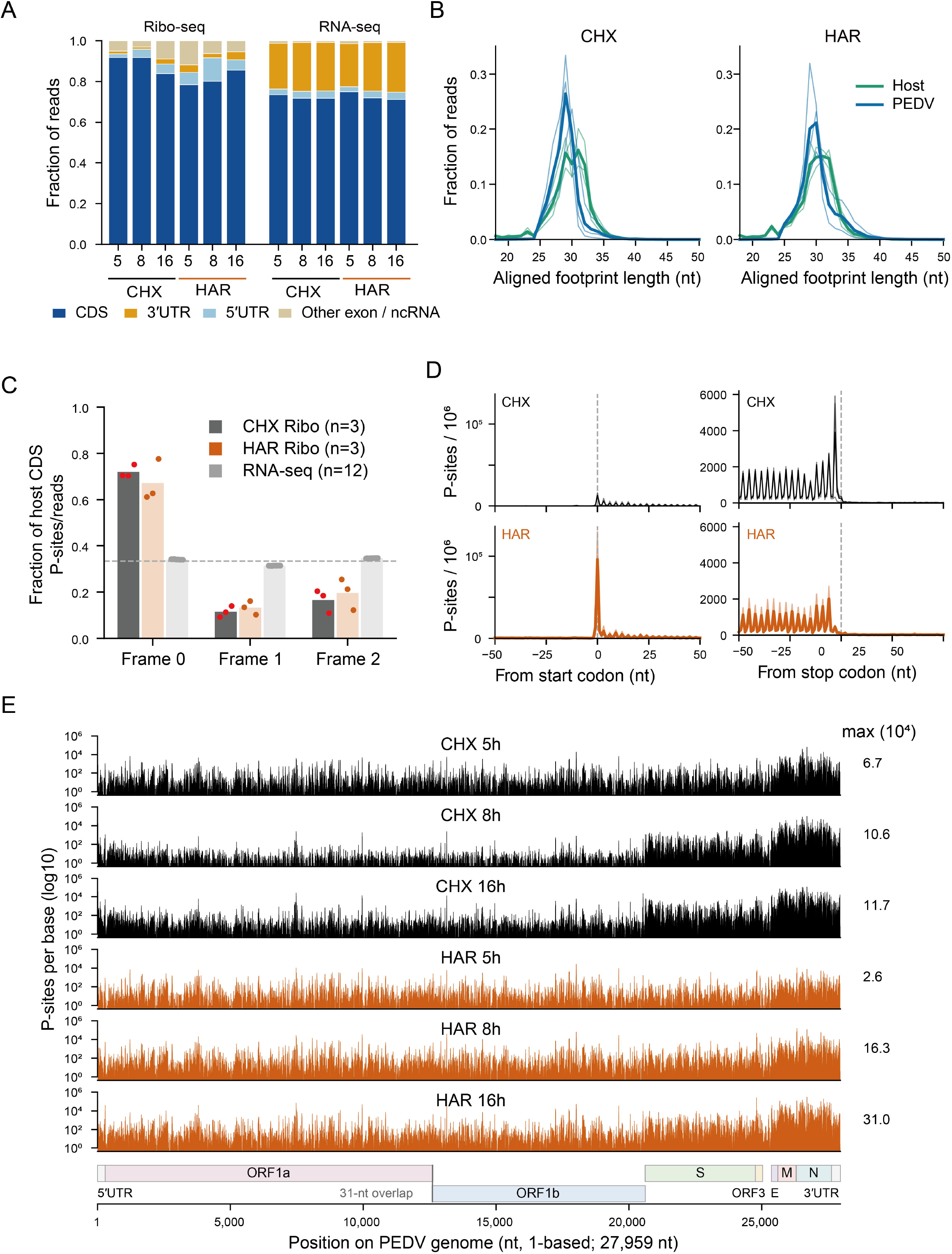
Ribosome profiling reveals characteristic translation signals and temporal redistribution of PEDV ribosome occupancy. (A) Distribution of P-sites across annotated transcript regions in Ribo-seq and RNA-seq libraries at 5, 8, and 16 hours post-infection (hpi). (B) Length distributions of host- and PEDV-derived ribosome-protected fragments (RPFs) in cycloheximide (CHX)- and harringtonine (HAR)-treated libraries. (C) Reading-frame distribution of host coding sequence (CDS) P-sites in CHX and HAR Ribo-seq libraries compared with RNA-seq. (D) Metagene profiles of host CDS P-sites around annotated start and stop codons under CHX and HAR treatment. Dashed lines indicate the reference codon positions. (E) Genome-wide PEDV P-site coverage at 5, 8, and 16 hpi under CHX and HAR treatment. Coverage is shown on a logarithmic scale, with the maximum P-site count indicated for each track. Viral genomic regions are annotated below.

Notably, reading-frame resolution depended strongly on RPF length. Analysis of host CDS-derived RPFs of different lengths revealed that, although fragments of 28 – 31 nt generally exhibited reading-frame preferences, their P-site distributions and background signals in alternative frames varied considerably (Fig S11). Among these, 29-nt RPFs showed the strongest enrichment in the annotated frame 0 across both CHX-and HAR-treated libraries, together with relatively low and balanced background signals in the +1 and +2 frames. We therefore preferentially used 29-nt RPFs for subsequent screening of overlapping ORF candidates to minimize asymmetric leakage of canonical reading-frame signals into alternative frames. This selection was particularly important for detecting weak translation-associated signals and comparing occupancy between the +1 and +2 reading frames.

Following quality assessment and P-site calibration, we examined the reading-frame occupancy patterns of PEDV. Compared with host CDSs, viral coding regions exhibited greater reading-frame heterogeneity, although this difference decreased as infection progressed (Fig S12). Among usable viral ORF P-sites, the fraction assigned to the annotated frame 0 increased from approximately 64% at 5 hpi to 72% at 16 hpi in CHX-treated libraries and from approximately 49% to 74% in HAR-treated libraries. Although host CDS-derived P-sites also exhibited a modest increase in frame-0 occupancy over time, the magnitude of this change was substantially smaller than that observed for PEDV (Fig S12A and S12B). In particular, the proportions of viral P-sites assigned to the +1 and +2 frames were considerably higher during early infection, especially in HAR-treated libraries, indicating greater alternative-frame occupancy at early time points. These patterns varied among viral genes. S, E, M, and N generally exhibited stronger enrichment in their annotated reading frames, particularly during later infection, whereas ORF1a and ORF1b showed comparatively lower and more variable frame-0 occupancy (Fig S12C).

Genome-wide P-site profiles further revealed pronounced spatial redistribution of viral ribosome occupancy during infection (Fig 4E). At 5 hpi, viral RPFs were broadly distributed across both the ORF1a/ORF1b replicase-coding regions and downstream structural protein-coding regions, with detectable ribosome occupancy in both genomic compartments. As infection progressed, particularly at 8 and 16 hpi, viral P-site signals became increasingly concentrated toward the genomic 3’ end, with especially prominent enrichment in the M and N coding regions. Signals associated with S and other structural protein-coding regions also increased. This pattern was observed under both CHX and HAR treatment and was consistent with the high accumulation of structural protein-associated viral RNA revealed by our RNA-seq coverage and sgRNA junction analyses. Given that ORF1a/ORF1b encode the replicase machinery required for viral RNA replication and transcription, whereas downstream structural proteins contribute to virion assembly, the temporal redistribution of ribosome occupancy is consistent with changing viral protein synthesis demands during infection.

### Results 5. Localized ribosome occupancy in PEDV untranslated regions exhibits reading-frame preferences

To investigate whether the untranslated regions (UTRs) of the PEDV genome harbor translation-associated activities beyond conventional coding annotations, we integrated cycloheximide (CHX)- and harringtonine (HAR)-treated Ribo-seq datasets to scan ribosomal P-site occupancy across the genomic 5’ and 3’ termini and compared localized enrichment signals with predicted short open reading frames (ORFs) (Fig 5). Although RNA-seq showed relatively continuous coverage across these regions, Ribo-seq signals were concentrated at several discrete positions, forming distinct local P-site peaks. Some of these peaks were reproducibly detected across multiple infection time points and under both drug treatments and exhibited reading-frame preferences relative to candidate ORFs, suggesting ribosome-associated occupancy that could not be explained solely by local RNA abundance.

**Fig 5.**
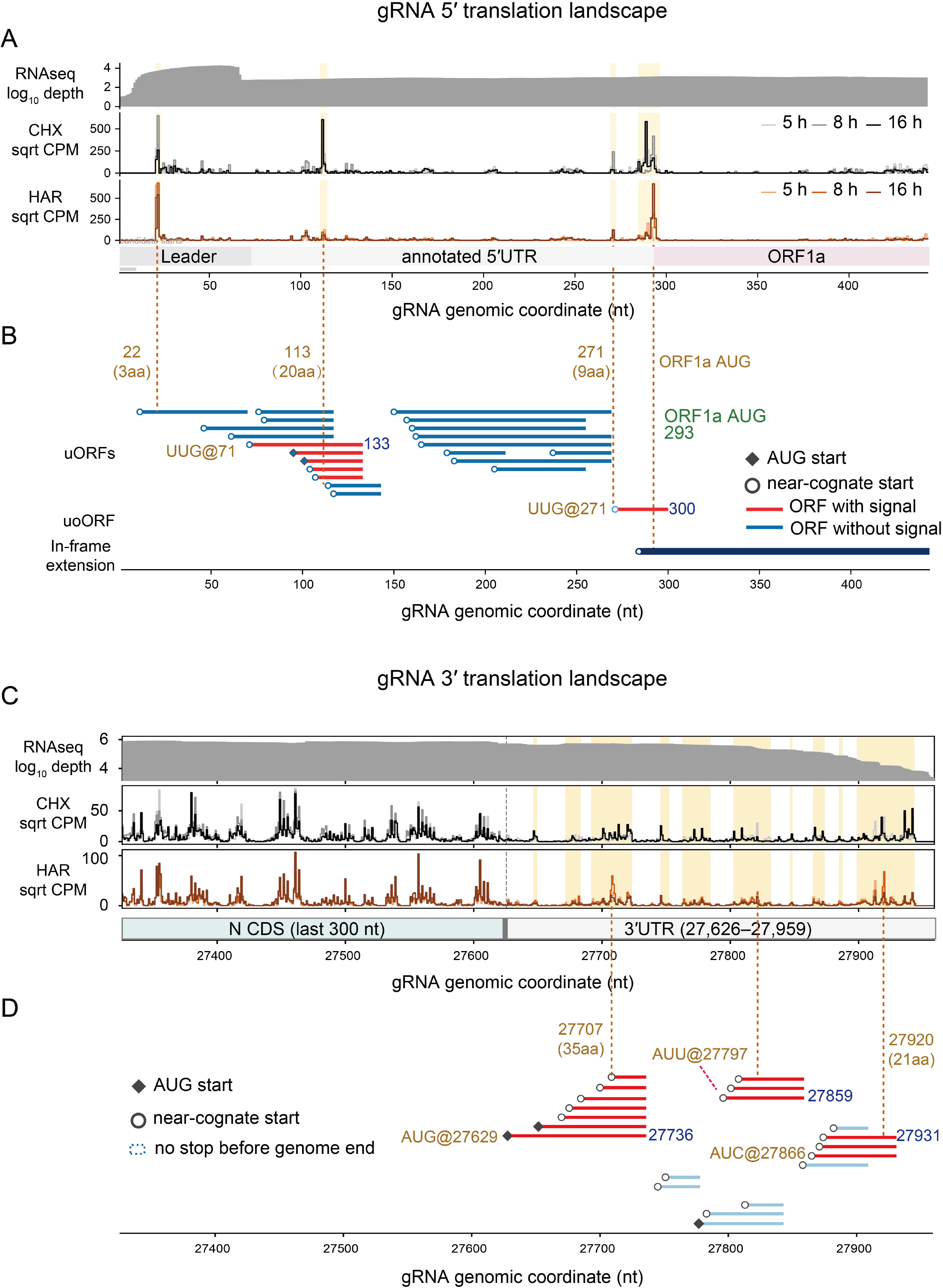
Localized ribosome occupancy identifies candidate translation-associated regions in PEDV untranslated regions. (A) RNA-seq coverage and CHX/HAR P-site profiles across the genomic 5’ region at 5, 8, and 16 hpi. Highlighted regions indicate major P-site hotspots. (B) Predicted upstream ORFs (uORFs), upstream overlapping ORFs, and a potential in-frame extension near the annotated ORF1a start codon. Red and blue lines distinguish candidate ORFs with and without associated local P-site signals, respectively. Filled diamonds indicate AUG starts and open circles indicate near-cognate starts. (C) RNA-seq coverage and CHX/HAR P-site profiles across the N gene terminus and genomic 3’ untranslated region (3’UTR). Highlighted regions indicate local P-site hotspots. (D) Predicted short ORFs in the 3’UTR, including candidates initiated at AUG or near-cognate codons. Colors distinguish candidates with and without local P-site support. Dashed terminal markers indicate ORFs without an in-frame stop codon before the genome end. P-site profiles in panels A and C are displayed as square-root-transformed counts per million (CPM). Genomic coordinates refer to the 27,959-nt PEDV reference genome.

At the genomic 5’ end, in addition to ribosome enrichment near the canonical ORF1a start codon (AUG at nucleotide 293), three major P-site hotspots were identified near nucleotides 22, 113, and 271 (Fig 5A and 5B). The signal near nucleotide 22 was particularly concentrated and exhibited pronounced candidate-frame P-site enrichment at all CHX and HAR time points. This region contained a predicted extremely short ORF encoding three amino acids. The hotspot near nucleotide 113 was located within a cluster of overlapping candidate short ORFs, including an approximately 20-amino-acid ORF defined by UUG at nucleotide 71 and extending to nucleotide 133. Recurrent CHX- and HAR-associated P-site signals were observed around nucleotides 103 – 113, with some P-sites consistent with the candidate reading frame. A distinct local enrichment signal was also detected near nucleotide 271, corresponding to an approximately 9-amino-acid candidate upstream overlapping ORF initiated at UUG at nucleotide 271 and extending to nucleotide 300, immediately upstream of and overlapping the canonical ORF1a initiation region. Frame-resolved local profiles further demonstrated that these hotspots were not uniformly distributed protected fragments across all three reading frames but instead displayed pronounced candidate-frame preferences at specific positions. Some hotspots also exhibited HAR-associated enrichment, supporting the possibility of genuine translation-related activity (Fig S13A).

At the genomic 3’ end, RNA-seq coverage extended from the terminal region of the N gene into the 3’UTR (nucleotides 27,626 – 27,959), whereas CHX and HAR profiles revealed several localized P-site hotspots near nucleotides 27,707, 27,820, and 27,919/27,920 (Fig 5C and 5D). The hotspot near nucleotide 27,707 was located within a cluster of candidate ORFs initiated at AUG at nucleotide 27,629, with the longest predicted product comprising approximately 35 amino acids. Reproducible ribosome-associated signals were detected at this position across infection time points. Notably, this candidate 3’UTR ORF was also predicted to be accessible from the noncanonical sgRNA generated by the 61→27608 leader–body junction identified in Fig S8B, linking a noncanonical transcript architecture to a potential translation-associated region in the 3’UTR. The hotspot near nucleotide 27,820 overlapped a candidate ORF region that included an approximately 20-amino-acid ORF extending from AUU at nucleotide 27,797 to nucleotide 27,859, and exhibited sharp local P-site enrichment in multiple CHX and HAR libraries. Finally, the hotspot near nucleotides 27,919/27,920 was adjacent to a cluster of candidate short ORFs, including one extending from AUC at nucleotide 27,866 to nucleotide 27,931. Strong local reading-frame preferences were observed in this region, with some signals being particularly prominent under HAR treatment (Fig S13B).

Collectively, these results reveal reproducible local P-site hotspots in both the 5’ and 3’UTRs of the PEDV genome, several of which overlap predicted short ORFs and exhibit candidate-frame preferences and HAR-associated enrichment. Although these signals do not establish that all corresponding short ORFs produce independent peptide products, they provide direct ribosome occupancy evidence for potential translation-associated activity in conventionally noncoding regions of the PEDV genome, suggesting that its translational landscape may extend beyond annotated protein-coding sequences.

### Results 6. The +1 overlapping reading frame of PEDV is enriched in potential novel coding genes supported by translational and proteomic evidence

To investigate whether PEDV may exploit alternative reading frames within canonical coding regions to produce previously unannotated proteins, we first examined the sequence composition of different reading frames in the context of viral overlapping gene evolution. Previous studies have demonstrated that RNA viruses can expand their limited coding capacity through overprinting, a process by which new protein-coding genes emerge within alternative reading frames of pre-existing coding sequences^14^. Because codons in alternative reading frames are formed by recombining nucleotides from adjacent codons in the canonical frame, codon usage bias, nucleotide composition at third codon positions, and constraints associated with stop-codon avoidance in existing genes can collectively influence the distribution of start and stop codons in alternative frames, thereby shaping their capacity to accommodate long open reading frames (ORFs). Previous investigations of viral overlapping genes have also suggested that, compared with the +2 frame, the +1 frame frequently contains more potential initiation codons and a more favorable distribution of termination codons, providing a sequence basis for the emergence of novel overlapping genes^15^.

Consistent with this pattern, an analysis of all available PEDV genome sequences from NCBI revealed pronounced sequence asymmetry among reading frames (S2 Table). Within canonical coding regions, AUG density in the +1 frame was approximately 18 sites per kilobase (kb), significantly higher than the approximately 7 sites per kb observed in the annotated frame 0 (*P* < 0.001), whereas stop-codon density in the +1 frame was only approximately 11 sites per kb. In contrast, AUG density in the +2 frame was approximately 1 site per kb, while stop-codon density reached approximately 43 sites per kb (Fig 6A). This pattern was consistently observed across all annotated PEDV ORFs (Fig S14A–G). These differences indicate that the +1 frame is more likely to satisfy the sequence requirements for both translation initiation and the formation of extended uninterrupted coding regions, whereas the +2 frame is strongly constrained by frequent termination codons. Systematic scanning of potential alternative ORFs further supported this distinction. We identified 68 ORFs longer than 40 amino acids (aa) in the +1 frame, compared with only three in the +2 frame (*P* < 0.001), and the maximum predicted ORF length was also greater in the +1 frame (Fig 6B). Thus, the PEDV genome contains a substantial potential overlapping coding space preferentially distributed in the +1 frame, consistent with patterns previously described for the emergence of overlapping genes in viruses.

**Fig 6.**
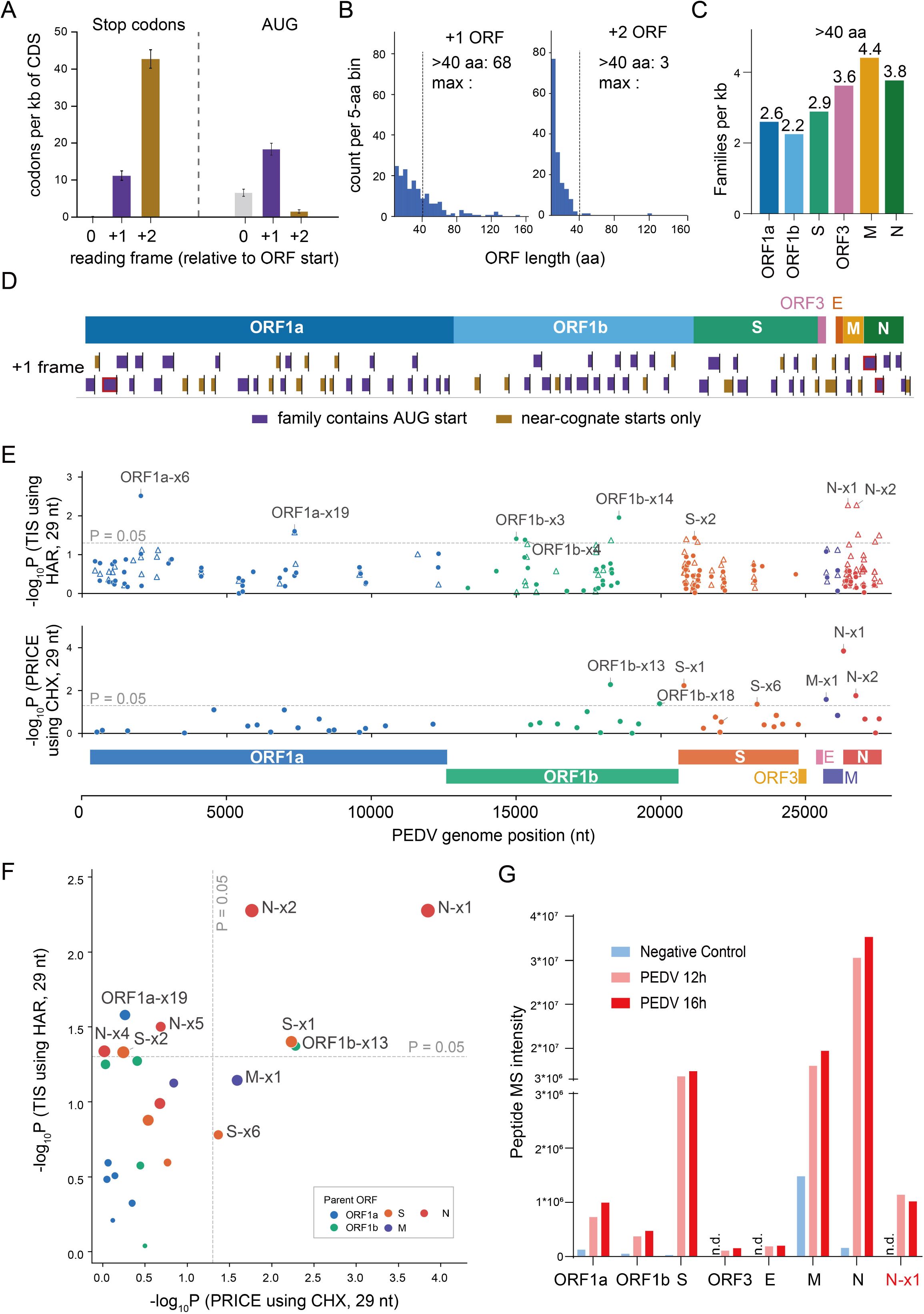
Alternative reading-frame analysis reveals extensive +1 overlapping coding potential and proteomic evidence for an unannotated PEDV peptide. (A) Densities of stop codons and AUG start codons in the annotated (0), +1, and +2 reading frames of PEDV coding sequences (CDSs). Values are expressed per kilobase of CDS. (B) Length distributions of predicted +1 and +2 overlapping ORFs. Dashed lines mark 40 amino acids (aa); the numbers of ORFs exceeding this threshold are indicated. (C) Density of +1 ORF families longer than 40 aa across individual annotated PEDV coding regions. (D) Genomic distribution of predicted +1 ORF families, distinguished by the presence of an AUG start codon or near-cognate starts only. (E) Genome-wide screening of candidate overlapping ORFs using HAR-based translation initiation-site analysis (HARTIS; upper panel) and CHX-based PRICE analysis (lower panel), restricted to 29-nt RPFs. Candidate positions are plotted against −log□□ P values; dashed lines indicate *P* = 0.05. Selected candidates are labeled. (F) Joint comparison of HAR-based initiation-associated enrichment and CHX-based PRICE evidence for candidate overlapping ORFs. Colors indicate the corresponding annotated parental ORFs. Dashed lines indicate *P* = 0.05. (G) Mass spectrometry peptide signal intensities for annotated PEDV proteins and the candidate N-x1 product in uninfected controls and PEDV-infected samples at 12 and 16 hpi. “n.d.” indicates not detected.

Beyond differences between reading frames, the distribution of long +1 ORFs was markedly heterogeneous across canonical PEDV genes. When normalized to coding sequence length, the M and N regions contained 4.4 and 3.8 families of +1 ORFs longer than 40 aa per kb, respectively, exceeding the corresponding densities of 2.6 and 2.2 in ORF1a and ORF1b (*P* < 0.001). The ORF3 and S regions exhibited intermediate densities of 3.6 and 2.9 ORF families per kb, respectively (Fig 6C). These candidates were distributed throughout the viral genome, with some containing AUG initiation codons and others possessing only near-cognate start codons (Fig 6D). Notably, the evolution of novel coding genes requires not only sequences capable of forming uninterrupted ORFs but also opportunities for those sequences to be expressed. Previous work on de novo gene birth and viral overprinting has proposed that low-level or occasional translation of previously unannotated sequences can generate initial peptide products, providing substrates for the emergence of new functions and subsequent natural selection, provided that these sequences occur in a translation-permissive environment^14,16^.

Accordingly, the translational environment surrounding alternative ORFs in different viral genes may influence the opportunities for novel overlapping proteins to emerge and persist. We therefore compared the density of long +1/+2 ORFs across canonical coding regions with RNA-normalized ribosome occupancy, expressed as translation efficiency (TE). Long overlapping ORF density was positively correlated with CHX-based TE (Spearman’s ρ = 0.83, *P* = 0.029, Fig S15A), whereas no significant association was observed with the corresponding HAR-based measure (ρ = 0.09, *P* = 0.46; Fig S15B). Together with the high abundance of canonical sgRNAs and the prominent ribosome occupancy of structural protein-coding regions described above, these observations highlight a potentially important convergence: the N and M regions not only harbor relatively abundant long overlapping ORFs but also reside within an expression environment characterized by abundant viral RNA templates and substantial ribosome occupancy during infection. Such regions may provide more opportunities for occasional alternative-frame translation to produce nascent peptide products, potentially increasing the likelihood that beneficial activities become accessible to natural selection and thereby creating conditions favorable for the emergence and evolution of novel viral proteins.

Because the coding potential of alternative ORFs may also be influenced by viral genetic background and evolutionary history, we next evaluated the conservation of initiation codon status and overall ORF architecture among selected overlapping ORF candidates using PEDV genomes from different genetic lineages (S16 Fig). The candidates exhibited distinct patterns of cross-lineage conservation. Several ORF1a-and N-associated candidates retained relatively high levels of ORF architectural integrity across multiple PEDV lineages. Among them, N-x1, a predicted 112-aa ORF representing the first potential alternative-frame ORF identified within the N coding region, showed broad conservation. In contrast, several S-associated candidates and N-x5 exhibited pronounced lineage-dependent variation in their initiation codon status or coding architecture. These results indicate that the potential overlapping coding space of PEDV does not consist exclusively of incidental ORFs present in a single reference genome. Instead, a subset of candidates possesses a relatively stable sequence basis across genetically divergent strains, supporting their prioritization for further investigation as potential viral expression products.

To distinguish candidate ORFs with translation-associated evidence from those predicted solely on the basis of sequence, we integrated CHX and HAR Ribo-seq datasets. Because canonical PEDV coding regions exhibited strong frame-0 ribosome occupancy, inaccuracies in P-site assignment could generate background signals in alternative reading frames. We therefore focused our primary analyses on 29-nt ribosome-protected fragments calibrated using annotated host coding sequences. As demonstrated by our preceding reading-frame assessment, this RPF length exhibited pronounced three-nucleotide periodicity and relatively low, balanced background leakage into the +1 and +2 frames, making it suitable for detecting comparatively weak alternative-frame signals within canonical coding regions. For CHX-treated datasets, we applied PRICE to evaluate ORF-level translation-associated signals^17^. For HAR-treated datasets, we assessed frame-specific local P-site enrichment around candidate initiation sites to identify translation initiation-associated signals. These complementary approaches provided evidence based on ribosome occupancy across candidate ORFs and localized enrichment near their predicted start sites^18^.

Systematic screening identified multiple potential +1 ORFs across the PEDV genome with varying degrees of translation-associated support (Fig 6E). Several candidates in the ORF1a, ORF1b, S, and N regions exhibited prominent HAR-associated enrichment near their predicted initiation sites, whereas CHX/PRICE analysis provided ORF-level support for an additional subset of candidates. Joint evaluation of the two datasets highlighted N-x1 and N-x2 as particularly noteworthy candidates (Fig 6F, Sup table 3). Both exhibited strong HAR initiation-associated signals, while N-x1 also showed prominent PRICE-based translation-associated evidence, providing complementary support from the two analytical approaches. Other candidates, including ORF1a-x19, ORF1b-x13, and S-x1, displayed varying degrees of translation-associated signals. Inspection of the underlying Ribo-seq profiles further revealed ribosome-protected fragments corresponding to the predicted alternative reading frames of N-x1, N-x2, ORF1b-x13, and S-x1, together with localized enrichment near the initiation regions of selected candidates (Fig S17A-D). These findings demonstrate that a subset of sequence-predicted overlapping ORFs in PEDV is associated with ribosome occupancy patterns consistent with candidate translation activity in infected cells, rather than being supported by sequence predictions alone.

Next, to determine whether these findings were cell-type-specific, we infected LLC-PK1 cells with PEDV GI-b (16 h) and performed RNA-seq, together with CHX- and HAR-treated Ribo-seq. The results showed strong correlations in P-site signal intensities between Vero and LLC-PK1 cells in both CHX- and HAR-treated Ribo-seq datasets (Spearman ρ = 0.86 and 0.92, respectively; Fig. S18A). Furthermore, the candidate ORFs N-x1, N-x2, ORF1b-x13, and S-x1 all exhibited prominent translation elongation- and initiation-associated signals in the corresponding +1 reading frame in LLC-PK1 cells (Fig. S18B). These results demonstrate that the translation-associated signals of these candidate ORFs are reproducible across different cell types.

Finally, we employed peptide mass spectrometry to investigate whether candidate overlapping ORFs could generate detectable protein products. By comparing uninfected controls with PEDV-infected samples collected at 12 and 16 hpi, we first observed marked increases in mass spectrometric signals corresponding to canonical viral proteins following infection (Fig. 6G and Fig. S19A-C). More importantly, peptide signals associated with the N-x1 candidate were detected at both 12 and 16 hpi but were absent from uninfected controls. Together with the translation-associated evidence obtained from HAR- and CHX-treated Ribo-seq datasets, these findings extend the evidence for N-x1 beyond alternative-frame sequence prediction and ribosome occupancy to the detection of a corresponding peptide product at the proteomic level.

Collectively, these results indicate that the potential overlapping coding capacity of PEDV is shaped by pronounced reading-frame asymmetry and gene-specific sequence features. The +1 frame provides a more favorable sequence environment for the formation of long alternative ORFs, while highly expressed structural protein-coding regions, particularly N and M, combine abundant overlapping ORF candidates with an active viral gene expression environment. This convergence may create opportunities for the emergence and evolutionary retention of novel overlapping proteins. The integration of ribosome profiling and peptide mass spectrometry further identifies N-x1 as a particularly promising candidate for a previously unannotated PEDV protein product.

## Discussion

The replication efficiency, cellular and tissue tropism, and pathogenic phenotypes of porcine epidemic diarrhea virus (PEDV) are closely associated with the regulation of viral RNA synthesis and protein expression. However, conventional genome annotations primarily focus on established viral proteins and may not fully capture the complex repertoire of viral RNAs and potential coding products generated during infection. By integrating time-resolved RNA sequencing (RNA-seq), cycloheximide (CHX)- and harringtonine (HAR)-treated ribosome profiling (Ribo-seq), comparative genomics, and peptide mass spectrometry, this study reveals multiple layers of complexity in PEDV RNA junction formation, ribosome occupancy, and alternative protein-coding potential. These findings provide new insights into how PEDV coordinates RNA replication and protein expression within the constraints of its compact genome.

First, we found that viral RNA accumulation across different genomic regions was not synchronized during PEDV infection. The pronounced relative enrichment of 3’-terminal sequences during early infection gradually decreased over time, whereas ribosome occupancy became increasingly concentrated in structural protein-coding regions at later stages. These changes may reflect coordinated regulation of viral RNA synthesis and protein production across infection stages. Establishing the nonstructural protein machinery required for viral RNA replication and transcription is essential, while continued structural protein synthesis is necessary for progeny virion assembly. The timing of these changes differed between MOI conditions, suggesting that the initial infection dose influences the temporal evolution of viral RNA composition and expression patterns. These findings provide a molecular perspective on PEDV infection kinetics and indicate that measurements of total viral RNA abundance alone may not adequately reflect the changing viral gene expression landscape during infection.

Second, discontinuous RNA junctions in PEDV were substantially more complex than predicted by the classical nested sgRNA model^5,8,9^. Although canonical sgRNAs accounted for the overwhelming majority of junction-supporting reads, noncanonical junctions exhibited extensive breakpoint diversity and a wide range of RNA architectures. Particularly notable was the overall declining gradient of 3’ breakpoint-supporting reads along the genome, interrupted by prominent enrichment peaks near canonical TRS-B sites. These observations led us to propose that low-probability template dissociation and rejoining may occur throughout negative-strand RNA synthesis^5–7^, whereas sequence features near canonical TRS-B sites may facilitate template switching and complementarity between the nascent negative-strand RNA and TRS-L may further favor joining to the leader region. The weaker TRS-L similarity associated with noncanonical leader–body junctions and the enrichment of short-sequence microhomology among different internal junction classes are consistent with this interpretation. Together, these findings suggest that canonical and noncanonical RNA junctions may share aspects of the underlying template-switching process while differing substantially in their preferred joining positions and sequence-dependent joining efficiencies.

The implications of noncanonical RNA junctions may extend beyond RNA architecture to viral protein-coding capacity. Some noncanonical leader–body RNAs were predicted to contain altered ORFs, including sequences potentially encoding N-terminally truncated proteins and alternative coding products generated by discontinuous joining. In addition, we identified localized ribosome occupancy signals in the viral 5’ and 3’ untranslated regions (UTRs), some of which displayed candidate-frame preferences and translation initiation-associated enrichment. These observations suggest that, beyond using canonical sgRNAs to express established viral proteins, PEDV may expand its repertoire of expression products through noncanonical RNA structures and alternative translation^8,9,11,12^. Of particular interest, certain noncanonical RNAs were predicted to encode truncated forms of the spike (S) protein. Given the central role of S in viral entry, host adaptation, and tissue tropism, determining whether these potential products are independently expressed and whether they influence viral protein localization, entry, or host interactions represents an important direction for future investigation.

A particularly important finding of this study is that the overlapping coding potential of PEDV exhibits pronounced reading-frame and gene-specific preferences. The codon composition of canonical coding regions favors the +1 frame by providing more potential initiation codons and fewer termination codons, thereby facilitating the formation of longer overlapping ORFs^14^. Moreover, structural protein-coding regions, particularly M and N, not only contain relatively high densities of long alternative ORFs but also reside in an expression environment characterized by abundant viral RNA and substantial ribosome occupancy. In the context of viral gene overprinting and de novo gene evolution, we propose that favorable alternative-frame sequence architectures and sufficient opportunities for expression may jointly promote the emergence and retention of novel overlapping viral proteins^14–16^. This concept is consistent with the occurrence of functional overlapping proteins within structural protein-coding regions of other coronaviruses^11,13^. By integrating complementary analyses of translation initiation-associated enrichment and ORF-level translation signals, we further prioritized several candidate overlapping ORFs. Importantly, the detection of infection-associated peptide mass spectrometry signals for N-x1 extends the evidence for hidden PEDV coding potential beyond sequence-based prediction to the proteomic detection of a candidate protein product.

These findings also raise important questions with virological and potential clinical relevance. Whether PEDV noncanonical RNAs and hidden proteins influence viral replication efficiency, cellular or tissue tropism, host immune responses, or strain-dependent differences in pathogenicity remains to be determined through functional studies^1,3,4,13^. Furthermore, the conservation and divergence of selected overlapping ORFs across PEDV lineages provide opportunities to investigate their evolutionary trajectories and potential associations with viral phenotypes. Future investigations involving genetically diverse PEDV strains, porcine intestinal epithelial cells, and in vivo infection models may help establish the biological functions of these noncanonical expression products.

Overall, this study demonstrates that the complexity of PEDV gene expression extends beyond the generation of canonical and noncanonical RNAs to include protein-coding products outside conventional genome annotations. Further characterization of this previously underexplored layer of viral gene expression may help identify molecular factors contributing to PEDV replication control, host adaptation, and pathogenesis.

## Materials and methods

### Cells and viruses

Porcine renal epithelial cells (LLC-PK1; ATCC CL-101) and African green monkey kidney cells (Vero; ATCC CCL-81) were maintained in Dulbecco’s modified Eagle’s medium (DMEM; Gibco, 11330032) supplemented with 10% fetal bovine serum (FBS; Cellmax, SA111.02) at 37°C in a humidified atmosphere containing 5% CO□. The PEDV XY2024 strain (GenBank accession number: [To be provided]), belonging to the GI-b lineage, was isolated and maintained in our laboratory. Viral stocks were propagated and titrated in Vero cells using DMEM supplemented with 5 μg/mL trypsin.

### Western blotting

Cells were lysed in RIPA lysis buffer (Solarbio, R0010) supplemented with protease inhibitors (Solarbio, R0100). Equal amounts of total protein were separated by SDS–polyacrylamide gel electrophoresis (SDS-PAGE) and transferred onto polyvinylidene difluoride (PVDF) membranes (Millipore, IPVH00010). After blocking with 5% skim milk, membranes were incubated overnight at 4°C with primary antibodies, followed by incubation with horseradish peroxidase (HRP)-conjugated secondary antibodies. Protein bands were detected using an enhanced chemiluminescence reagent (Oriscience, PD202). A laboratory-generated antibody against the PEDV nucleocapsid (N) protein was used at a dilution of 1:500.

### Quantification of viral RNA by RT-qPCR

Total RNA was extracted from mock-infected and PEDV-infected Vero cells using TRIzol reagent (Invitrogen). Complementary DNA (cDNA) was synthesized using a reverse transcription kit (Vazyme, R312-01). Quantitative real-time PCR was performed using an iQ5 real-time PCR detection system (Bio-Rad) with SYBR Green-based detection and PEDV-specific primers. Relative PEDV RNA levels were normalized to β-actin expression.

### RNA-seq and Ribosome profiling library preparation

Vero cells were infected with PEDV at multiplicities of infection (MOIs) of 0.1 and 1 and collected at 3, 5, 8, 12, and 16 hours post-infection (hpi) for RNA-seq analysis. The principal Ribo-seq experiment was conducted using PEDV-infected Vero cells collected at 5, 8, and 16 hpi. RNA-seq and ribosome profiling libraries were prepared by Gene Denovo Biotechnology Co., Ltd. (Guangzhou, China) as described previously^18–20^.

For cycloheximide (CHX)-treated samples, infected cells were exposed to CHX for 5 min before collection. For harringtonine (HAR)-treated samples, cells were first exposed to HAR for 3 min, followed by CHX treatment for an additional 5 min. These treatments were used to obtain complementary profiles of elongation-associated ribosome occupancy and translation initiation-associated enrichment. For ribosome profiling, pretreated cells were lysed, and the resulting lysates were subjected to nuclease digestion using RNase I (New England Biolabs) and DNase I (New England Biolabs; 6 μL) for 45 min at room temperature with gentle mixing. Ribosome-protected fragments (RPFs) were subsequently recovered and used for library construction. RNA-seq libraries were sequenced using paired-end 150-bp sequencing. Sequencing was performed on an Illumina HiSeq X Ten platform, according to the sequencing information provided by the library preparation service. For Ribo-seq analyses, the first sequencing read (R1) was used as the input for subsequent processing. The principal RNA-seq dataset comprised 20 libraries representing two MOIs, two drug-treatment conditions, and five infection time points, with one library per experimental condition. Six Ribo-seq libraries represented the CHX and HAR treatments at 5, 8, and 16 hpi. These sequencing datasets did not include independent biological replicates for each condition.

### RNA-seq preprocessing and reference alignment

Raw paired-end RNA-seq reads were subjected to adapter removal and quality filtering using fastp^19^. Filtered reads were aligned with STAR^21^ (version 2.7.11b) against a combined reference containing the *Chlorocebus sabaeus* genome assembly ChlSab1.1 and the PEDV GI-b reference genome (27,959 nucleotides). To accommodate discontinuous viral RNA alignments, STAR was configured using splice-permissive settings, including a maximum intron length of 30,000 nt, a maximum mate gap of 30,000 nt, a maximum of six mismatches per read, a mismatch fraction threshold of 0.08, and a maximum of 10 reported mapping loci. The minimum splice-junction overhang was set to 8 nt. Noncanonical splice-junction penalties were adjusted to permit the detection of discontinuous viral alignments. Aligned reads were sorted and indexed using SAMtools^22^. Uniquely mapped alignments were identified using a mapping quality value of 255 and the alignment tag NH:i:1. Host- and virus-derived alignments were distinguished according to their reference contigs. The primary viral RNA coverage analyses were performed using non-deduplicated alignments. Because the libraries did not contain unique molecular identifiers, coordinate-based duplicate removal was not used for the main analyses. Separately, duplicate-marked and duplicate-removed BAM files were generated using the SAMtools fixmate and markdup procedures for sensitivity analyses.

Public PEDV RNA-seq datasets were retrieved from the European Nucleotide Archive (ENA), including PRJNA730044 (HX- and GDS01-infected IPEC-J2 cells), PRJNA1069035 (LJX-infected piglet jejunum), and PRJNA1203018 (GIIb infected LLC-PK cells)^23,24^. As complete reference genomes were unavailable for LJX and GIIa P10, strain-specific reference genomes were assembled de novo and polished using sequencing reads. The corresponding viral reads were mapped to the respective reference genomes using minimap2. For cross-strain comparisons, assembled genomes were aligned to the PEDV GI-b reference to convert junction coordinates into a common genomic coordinate system.

### Analysis of viral RNA coverage and the 3’/5’ coverage ratio

Per-nucleotide sequencing depth across the PEDV genome was calculated using SAMtools depth from uniquely mapped, non-deduplicated alignments. To characterize regional differences in viral RNA accumulation, a 3’/5’ coverage ratio was calculated separately for each RNA-seq library. The 3’ region was defined as nucleotides 20,610–27,625, corresponding approximately to the S-to-N genomic interval. The 5’ region was defined as nucleotides 1,101–20,613, spanning the principal replicase-coding region.

The ratio was calculated as the mean nucleotide coverage of the defined 3’ region divided by the mean nucleotide coverage of the defined 5’ region:

3’/5’ coverage ratio = Mean coverage of the 3’ region / Mean coverage of the 5’ region.

This measure represents relative sequencing coverage between two genomic regions rather than the absolute molecular abundance ratio of sgRNAs to genomic RNA, because sequences within the 3’ interval are shared by genomic RNA and multiple nested sgRNAs. Mean regional coverage was normalized to the total number of mapped host and viral fragments per million. Within each MOI and infection time point, the normalized values from CHX- and HAR-associated RNA-seq libraries were averaged.

### Identification and classification of PEDV RNA junctions

To identify discontinuous viral RNA junctions, adapter-trimmed RNA-seq reads were independently aligned to the PEDV reference genome using minimap2 in splice alignment mode, with the parameters -ax splice -un -G 30000 -k 13 -w 5 --secondary=no. Trimmed R1 and R2 sequences were processed as individual alignment records. Candidate junctions were extracted from skipped alignment regions indicated by the CIGAR operation N, with a minimum junction gap of 50 nt. Junctions were retained when they fulfilled the following criteria: mapping quality ≥20, at least 15 nt of aligned sequence on each side of the junction, and a reliable primary alignment. When both mates from the same sequencing fragment supported an identical junction, the fragment was counted only once. PCR duplicate removal was not applied.

Junctions were classified into four categories according to leader involvement and genomic separation:

TRS-L-dependent canonical junctions: junctions with donor positions within the defined leader-associated window and acceptor positions within ±5 nt of the canonical S (20,597), ORF3 (24,718), E (25,342), M (25,597), or N (26,285) acceptor site. TRS-L-dependent noncanonical junctions: junctions involving a leader-associated donor position but an acceptor outside the canonical windows. TRS-L-independent distant junctions: junctions without a leader-associated donor and with a genomic separation of at least 1,000 nt. TRS-L-independent local junctions: junctions without a leader-associated donor and with a genomic separation below 1,000 nt.

### Prediction of coding consequences of noncanonical RNA junctions

The potential coding consequences of noncanonical junctions were assessed relative to annotated PEDV coding sequences. The first downstream AUG within the acceptor-associated coding region was identified and classified according to its reading frame relative to the canonical ORF. In parallel, hypothetical reading-frame continuity across each junction was evaluated independently. These analyses were used to distinguish potential canonical-frame, alternative-frame, and noncoding junction configurations.

### TRS-L sequence similarity and junction microhomology analysis

TRS-L similarity was assessed using the 12-nt leader reference sequence (5’-CAACTAAACGAA-3’; nt 64–75). For each junction acceptor, the maximum nucleotide identity across 41 ungapped alignments (offsets −20 to +20 nt) was calculated and compared with a background generated from 200 dinucleotide-preserving shuffles of the acceptor sequence context. Junction microhomology was defined as the number of consecutive identical nucleotides flanking donor and acceptor breakpoints in the positive-strand reference sequence. Background distributions were generated through 10,000 donor–acceptor permutations within each junction class, stratified by genomic-span quartiles. The five dominant canonical junctions were evaluated using position-based backgrounds. Microhomology enrichment was calculated as the difference between observed and background mean values, with each unique junction treated as an independent observation.

### Prediction of RNA pairing free energy

RNA pairing stability was evaluated using RNAduplex (ViennaRNA) and 30-nt sequences surrounding each junction breakpoint^25^. Two configurations were examined: pairing between the reverse-complemented donor sequence and the acceptor sequence, representing potential nascent negative-strand re-pairing, and direct pairing between the two positive-strand sequences. Minimum free energy (MFE) differences were calculated relative to span-stratified donor–acceptor permutation backgrounds. Statistical significance was assessed using 10,000 two-sided empirical permutations with Holm correction. Bootstrap confidence intervals and Spearman correlations were used to assess effect estimates and relationships with junction abundance or recurrence, respectively.

### Identification of translation-associated signals in PEDV untranslated regions

Ribosome occupancy in the PEDV 5’ and 3’ untranslated regions (UTRs) was examined using calibrated P-site distributions from CHX- and HAR-treated Ribo-seq libraries. For the genomic 5’ region, positive-strand P-site occupancy was examined across nucleotides 1–443. For the 3’UTR, nucleotides 27,626–27,959 were examined separately from the upstream N coding sequence. Local P-site peaks were identified using a minimum of 10 P-sites at a candidate position and a signal at least fivefold greater than the median occupancy within a surrounding ±50-nt window. A hotspot was considered recurrent when a peak was detected within ±1 nt of the same position in at least two libraries under the same drug treatment. Potential short ORFs were enumerated on the positive strand using AUG and seven near-cognate initiation codons (CUG, GUG, UUG, AUA, AUU, AUC, and ACG). Candidate ORFs were extended to the first in-frame termination codon, and an 8-aa minimum length was used in the computational candidate enumeration. HAR-associated initiation enrichment in the 3’UTR was additionally examined by comparing P-site density near candidate start sites with local flanking occupancy. Candidate-frame preferences were evaluated relative to the reading frame of each predicted UTR ORF. The 5’UTR and 3’UTR analyses employed separately calibrated RPF-length subsets. Therefore, their local occupancy measurements were evaluated within their respective analytical frameworks rather than assuming that an identical footprint-length filter was applied to both regions.

### PEDV genome collection and sequence curation

Complete PEDV genome sequences were retrieved from the NCBI Nucleotide database using the Entrez E-utilities on September 24, 2026^26^. Searches targeted records annotated as porcine epidemic diarrhea virus and containing complete genomic sequences. Downloaded records were screened to retain sequences between 25,000 and 34,000 nt in length, with no more than 1% ambiguous nucleotides. Incomplete genomic fragments, synthetic constructs, patent-associated sequences, cloning vectors, and records without a reliable viral assignment were excluded. After sequence filtering and removal of identical sequences, 1,041 publicly retrieved PEDV genomes were retained. These were merged with an existing local PEDV genome collection, and duplicate sequences were removed to obtain a final dataset comprising 1,083 genomes. Available lineage annotations were collected and standardized where possible. For genomes without a verified lineage assignment, sequence similarity to reference genomes was examined to support provisional classification. The distinction between verified and computationally inferred lineage assignments was retained during downstream analysis.

### Analysis of alternative-frame start and stop codon distributions

The sequence composition of alternative reading frames was examined relative to annotated PEDV protein-coding sequences. For each canonical coding region, the annotated reading frame was designated frame 0, and the two alternative reading frames were designated +1 and +2 according to their nucleotide offsets relative to the parent coding sequence. The positions of AUG codons and the three canonical termination codons (UAA, UAG, and UGA) were enumerated within each reading frame. Start- and stop-codon densities were normalized to the number of nucleotides analyzed and expressed as the number of sites per kilobase. Frame-specific densities were compared to characterize differences in the sequence capacity of alternative frames to support extended uninterrupted coding intervals.

### Prediction of overlapping open reading frames

Candidate overlapping ORFs were identified within annotated PEDV coding regions by scanning alternative reading frames for eligible initiation codons followed by uninterrupted translation to an in-frame termination codon. The eligible initiation codons were AUG, CUG, GUG, UUG, AUA, AUU, AUC, and ACG. Candidate ORFs terminated at the first downstream in-frame UAA, UAG, or UGA codon. Predicted peptide length excluded the termination codon. For the principal overlapping ORF analysis, candidate sequences were required to encode more than 40 amino acids. Multiple candidate initiation sites sharing the same termination codon within a given parent coding region and reading frame were grouped into a single stop-anchored ORF family. Individual initiation sites within a family were retained as alternative candidate isoforms. Candidate ORFs were assigned to their respective parent canonical coding regions, including ORF1a, ORF1b, S, ORF3, E, M, and N. Family densities were normalized to the length of the corresponding annotated coding region and expressed as the number of ORF families per kilobase. A separate genome-wide stop-anchored scanning procedure was used for cross-strain sequence analysis. In this analysis, eligible initiation sites were enumerated across all three genomic frames and grouped according to shared in-frame termination positions.

### HAR-based analysis of translation initiation-associated signals

Translation initiation-associated ribosome occupancy in candidate overlapping ORFs was evaluated using HAR-treated Ribo-seq datasets and a custom frame-resolved framework (HARTIS). P-site counts were assigned to the canonical frame (n□), candidate alternative frame (n□), and remaining alternative frame (n□). To distinguish candidate-frame enrichment from background frame leakage, the analysis was conditioned on the total alternative-frame occupancy, m = n□ + n□, rather than directly comparing n□ with n□.

A reference set of 15,281 high-confidence host coding sequences was used to estimate expected alternative-frame occupancy from 256 sequence-context categories with cross-fitting. Candidate-frame enrichment was quantified using a depth-standardized residual:

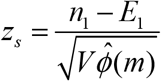

where E□ and V denote the context-dependent expected candidate-frame count and its variance, respectively, and φ□(m) represents the depth-dependent dispersion correction. Host reference windows were matched by alternative-frame depth (log□m) and canonical-frame P-site density. The empirical upper-tail probability was calculated as:

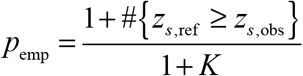

where K is the number of matched host reference windows. Analyses were performed separately for each library, RPF length, and initiation window (W, the start-codon position).

For viral regions exceeding the calibrated host-depth range, a capped-reference procedure was applied using the highest supported host-depth interval. The context-adjusted candidate-frame fraction was defined as:

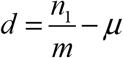

where μ is the expected candidate-frame fraction estimated from host sequence contexts. The corresponding capped-reference probability was:

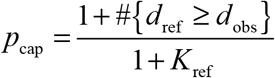

Analyses were performed separately for each HAR-treated library using exclusively 29-nt ribosome-protected fragments (RPFs), with the W1 initiation window defined as the single P-site position corresponding to the candidate start codon. For Figure 6, the minimum available W1 capped-reference probability across candidate starts, HAR libraries, and eligible RPF lengths was used for visualization without multiple-testing correction. These capped-reference probabilities were treated as exploratory because viral occupancy exceeded the validated host-reference range.

### CHX-based analysis of alternative ORF translation using PRICE

CHX-treated Ribo-seq data were independently analyzed using PRICE, implemented in the Gedi software framework, to assess ORF-level translation-associated signals. The primary PRICE^17^ analysis was restricted to 29-nt footprints. Host-derived alignments were used to establish the translation model, which was subsequently applied to PEDV-derived alignments using the corresponding viral genome annotation. Candidate results were compared with the independently enumerated alternative ORF families using their initiation and termination coordinates. For integration with HAR-based initiation analysis, raw PRICE probabilities were summarized across the three selected CHX libraries using the minimum observed value for each candidate. HAR-based capped-reference probabilities were summarized separately across eligible candidate initiation sites, libraries, and calibrated footprint lengths.

### Analysis of start and stop codon distributions across reading frames

To characterize the coding potential of alternative reading frames, the PEDV positive-strand genome was analyzed relative to the annotated viral coding sequences. For each canonical ORF, the annotated reading frame was designated frame 0, and the two alternative frames were designated +1 and +2. AUG initiation codons and canonical termination codons (UAA, UAG, and UGA) were counted separately in each frame. Codon densities were calculated as the number of corresponding codons divided by the analyzed sequence length and expressed per kilobase. Frame-specific codon distributions were compared to evaluate the sequence constraints associated with the formation of overlapping ORFs.

### Parallel reaction monitoring (PRM) mass spectrometry

Targeted proteomic analysis was performed using parallel reaction monitoring (PRM) mass spectrometry to detect candidate PEDV-derived peptides in infected Vero cells collected at 12 and 16 hpi, with uninfected cells serving as controls^27^. Protein samples were subjected to tryptic digestion overnight at an enzyme-to-protein ratio of 1:50, followed by additional digestion at a ratio of 1:100 for 4 h. The resulting peptides were desalted using C18 solid-phase extraction and analyzed by liquid chromatography–tandem mass spectrometry (LC–MS/MS) using a timsTOF Pro mass spectrometer. Targeted peptide signals were evaluated based on their chromatographic profiles and fragment-ion patterns.

### Cross-lineage conservation of candidate overlapping ORFs

To evaluate the preservation of candidate overlapping ORFs across genetically diverse PEDV genomes, reference-based candidate ORF coordinates were mapped to the curated genome collection. For each candidate ORF, the mapped initiation and termination regions were examined to determine whether the relevant coding architecture was preserved. Initiation codons were categorized according to the retention of the original codon, substitution by another eligible initiation codon, or loss of an eligible initiation codon. At the ORF-family level, a candidate was considered retained when the corresponding termination position and reading frame remained intact and at least one eligible initiation codon could generate an uninterrupted coding interval meeting the specified length criterion. Genomes containing incomplete sequence coverage, ambiguous nucleotides, or unresolved coding boundaries that prevented reliable assessment were classified as not assessable and excluded from the corresponding retention-rate denominator.

## Statistical analysis

Statistical analyses were conducted according to the characteristics of each dataset. For junction-based sequence analyses, unique donor–acceptor pairs were treated as the primary analytical units rather than weighting individual junctions by sequencing read abundance. Randomized background distributions were generated using sequence shuffling or donor–acceptor pair permutation, as appropriate for the tested sequence feature. Empirical two-sided permutation probabilities were calculated using 10,000 iterations and corrected using the Holm procedure for predefined comparisons among noncanonical junction classes. Confidence intervals for sequence-feature enrichment were calculated using bootstrap resampling. Spearman rank correlation coefficients were used to assess monotonic relationships between junction-associated sequence features and recurrence or supporting read abundance.

## Acknowledgments

Data and code availability

All high-throughput sequencing data are available from NCBI under accession: PRJNA1554149. Scripts for performing all analyses and generating figures in this manuscript are available at https://github.com/guanguiwensy/PEDV_RNAseq_Riboseq

## Author contributions

Conceptualization: Guiwen Guan, Yong Huang.

Data curation: Guiwen Guan, Ran Zhang.

Funding acquisition: Yong Huang, Guiwen Guan

Investigation: Guiwen Guan, Ran Zhang, Jin Wang, Hanwen Zhang, Leyu Hu, Mingjun Zhu, Qian Du, Lu Han, Xin Ran

Project administration: Yong Huang

Writing – original draft: Guiwen Guan, Yong Huang, Ran Zhang Writing – review & editing: Yong Huang, Qian Du

## References

1. Jung, K., Saif, L.J., and Wang, Q. (2020). Porcine epidemic diarrhea virus (PEDV): An update on etiology, transmission, pathogenesis, and prevention and control. Virus Research 286, 198045. 10.1016/j.virusres.2020.198045.

2. Wei, M.-Z., Chen, L., Zhang, R., Chen, Z., Shen, Y.-J., Zhou, B.-J., Wang, K.-G., Shan, C.-L., Zhu, E.-P., and Cheng, Z.-T. (2024). Overview of the recent advances in porcine epidemic diarrhea vaccines. The Veterinary Journal 304, 106097. 10.1016/j.tvjl.2024.106097.

3. Li, Z., Ma, Z., Dong, L., Yang, T., Li, Y., Jiao, D., Han, W., Zheng, H., and Xiao, S. (2022). Molecular Mechanism of Porcine Epidemic Diarrhea Virus Cell Tropism. mBio 13, e03739–03721. 10.1128/mbio.03739-21.

4. Li, W., Hangalapura, B.N., van den Elzen, P., van den Born, E., van Kuppeveld, F.J.M., Rottier, P.J.M., and Bosch, B.-J. (2025). Spike gene variability in porcine epidemic diarrhea virus as a determinant for virulence. Journal of Virology 99, e02165–02124. 10.1128/jvi.02165-24.

5. Sola, I., Almazán, F., Zúñiga, S., and Enjuanes, L. (2015). Continuous and Discontinuous RNA Synthesis in Coronaviruses. Annual Review of Virology 2, 265–288. 10.1146/annurev-virology-100114-055218.

6. Zúñiga, S., Sola, I., Alonso, S., and Enjuanes, L. (2004). Sequence motifs involved in the regulation of discontinuous coronavirus subgenomic RNA synthesis. Journal of Virology 78, 980–994. 10.1128/JVI.78.2.980-994.2004.

7. Mateos-Gomez, P.A., Morales, L., Zúñiga, S., Enjuanes, L., and Sola, I. (2013). Long-distance RNA-RNA interactions in the coronavirus genome form high-order structures promoting discontinuous RNA synthesis during transcription. Journal of Virology 87, 177–186. 10.1128/JVI.01782-12.

8. Kim, D., Lee, J.-Y., Yang, J.-S., Kim, J.W., Kim, V.N., and Chang, H. (2020). The Architecture of SARS-CoV-2 Transcriptome. Cell 181, 914–921.e910. 10.1016/j.cell.2020.04.011.

9. Nomburg, J., Meyerson, M., and DeCaprio, J.A. (2020). Pervasive generation of non-canonical subgenomic RNAs by SARS-CoV-2. Genome Medicine 12, 108. 10.1186/s13073-020-00802-w.

10. Ingolia, N.T., Ghaemmaghami, S., Newman, J.R.S., and Weissman, J.S. (2009). Genome-wide analysis in vivo of translation with nucleotide resolution using ribosome profiling. Science 324, 218–223. 10.1126/science.1168978.

11. Finkel, Y., Mizrahi, O., Nachshon, A., Weingarten-Gabbay, S., Morgenstern, D., Yahalom-Ronen, Y., Tamir, H., Achdout, H., Stein, D., Israeli, O., et al. (2021). The coding capacity of SARS-CoV-2. Nature 589, 125–130. 10.1038/s41586-020-2739-1.

12. Kim, D., Kim, S., Park, J., Chang, H.R., Chang, J., Ahn, J., Park, H., Park, J., Son, N., Kang, G., et al. (2021). A high-resolution temporal atlas of the SARS-CoV-2 translatome and transcriptome. Nature Communications 12, 5120. 10.1038/s41467-021-25361-5.

13. Weingarten-Gabbay, S., Klaeger, S., Sarkizova, S., Pearlman, L.R., Chen, D.-Y., Gallagher, K.M.E., Bauer, M.R., Taylor, H.B., Dunn, W.A., Tarr, C., et al. (2021). Profiling SARS-CoV-2 HLA-I peptidome reveals T cell epitopes from out-of-frame ORFs. Cell 184, 3962–3980.e3917. 10.1016/j.cell.2021.05.046.

14. Sabath, N., Wagner, A., and Karlin, D. (2012). Evolution of viral proteins originated de novo by overprinting. Molecular Biology and Evolution 29, 3767–3780. 10.1093/molbev/mss179.

15. Willis, S., and Masel, J. (2018). Gene Birth Contributes to Structural Disorder Encoded by Overlapping Genes. Genetics 210, 303–313. 10.1534/genetics.118.301249.

16. Carvunis, A.-R., Rolland, T., Wapinski, I., Calderwood, M.A., Yildirim, M.A., Simonis, N., Charloteaux, B., Hidalgo, C.A., Barbette, J., Santhanam, B., et al. (2012). Proto-genes and de novo gene birth. Nature 487, 370–374. 10.1038/nature11184.

17. Erhard, F., Halenius, A., Zimmermann, C., L’Hernault, A., Kowalewski, D.J., Weekes, M.P., Stevanovic, S., Zimmer, R., and Dölken, L. (2018). Improved Ribo-seq enables identification of cryptic translation events. Nature Methods 15, 363–366. 10.1038/nmeth.4631.

18. Ingolia, N.T., Lareau, L.F., and Weissman, J.S. (2011). Ribosome profiling of mouse embryonic stem cells reveals the complexity and dynamics of mammalian proteomes. Cell 147, 789–802. 10.1016/j.cell.2011.10.002.

19. Chen, S., Zhou, Y., Chen, Y., and Gu, J. (2018). fastp: an ultra-fast all-in-one FASTQ preprocessor. Bioinformatics 34, i884–i890. 10.1093/bioinformatics/bty560 PMID - 30423086.

20. McGlincy, N.J., and Ingolia, N.T. (2017). Transcriptome-wide measurement of translation by ribosome profiling. Methods 126, 112–129. 10.1016/j.ymeth.2017.05.028.

21. Dobin, A., Davis, C.A., Schlesinger, F., Drenkow, J., Zaleski, C., Jha, S., Batut, P., Chaisson, M., and Gingeras, T.R. (2013). STAR: ultrafast universal RNA-seq aligner. Bioinformatics 29, 15–21. 10.1093/bioinformatics/bts635 PMID - 23104886.

22. Li, H., Handsaker, B., Wysoker, A., Fennell, T., Ruan, J., Homer, N., Marth, G., Abecasis, G., Durbin, R., and Genome Project Data Processing, S. (2009). The Sequence Alignment/Map format and SAMtools. Bioinformatics 25, 2078–2079. 10.1093/bioinformatics/btp352 PMID - 19505943.

23. Leinonen, R., Sugawara, H., Shumway, M., and International Nucleotide Sequence Database, C. (2011). The sequence read archive. Nucleic Acids Res 39, D19–21. 10.1093/nar/gkq1019.

24. Barrett, T., Wilhite, S.E., Ledoux, P., Evangelista, C., Kim, I.F., Tomashevsky, M., Marshall, K.A., Phillippy, K.H., Sherman, P.M., Holko, M., et al. (2013). NCBI GEO: archive for functional genomics data sets--update. Nucleic Acids Res 41, D991–995. 10.1093/nar/gks1193.

25. Lorenz, R., Bernhart, S.H., Hoener zu Siederdissen, C., Tafer, H., Flamm, C., Stadler, P.F., and Hofacker, I.L. (2011). ViennaRNA Package 2.0. Algorithms for Molecular Biology 6, 26. 10.1186/1748-7188-6-26 PMID - 22115189.

26. Sayers, E.W., Bolton, E.E., Brister, J.R., Canese, K., Chan, J., Comeau, D.C., Connor, R., Funk, K., Kelly, C., Kim, S., et al. (2022). Database resources of the national center for biotechnology information. Nucleic Acids Res 50, D20–D26. 10.1093/nar/gkab1112.

27. Peterson, A.C., Russell, J.D., Bailey, D.J., Westphall, M.S., and Coon, J.J. (2012). Parallel reaction monitoring for high resolution and high mass accuracy quantitative, targeted proteomics. Mol Cell Proteomics 11, 1475–1488. 10.1074/mcp.O112.020131.

