## Supplementary figure for "Integrated Transcriptomic and Translatomic Profiling Reveals Noncanonical RNA Junctions and Hidden Coding Potential in Porcine Epidemic Diarrhea Virus"

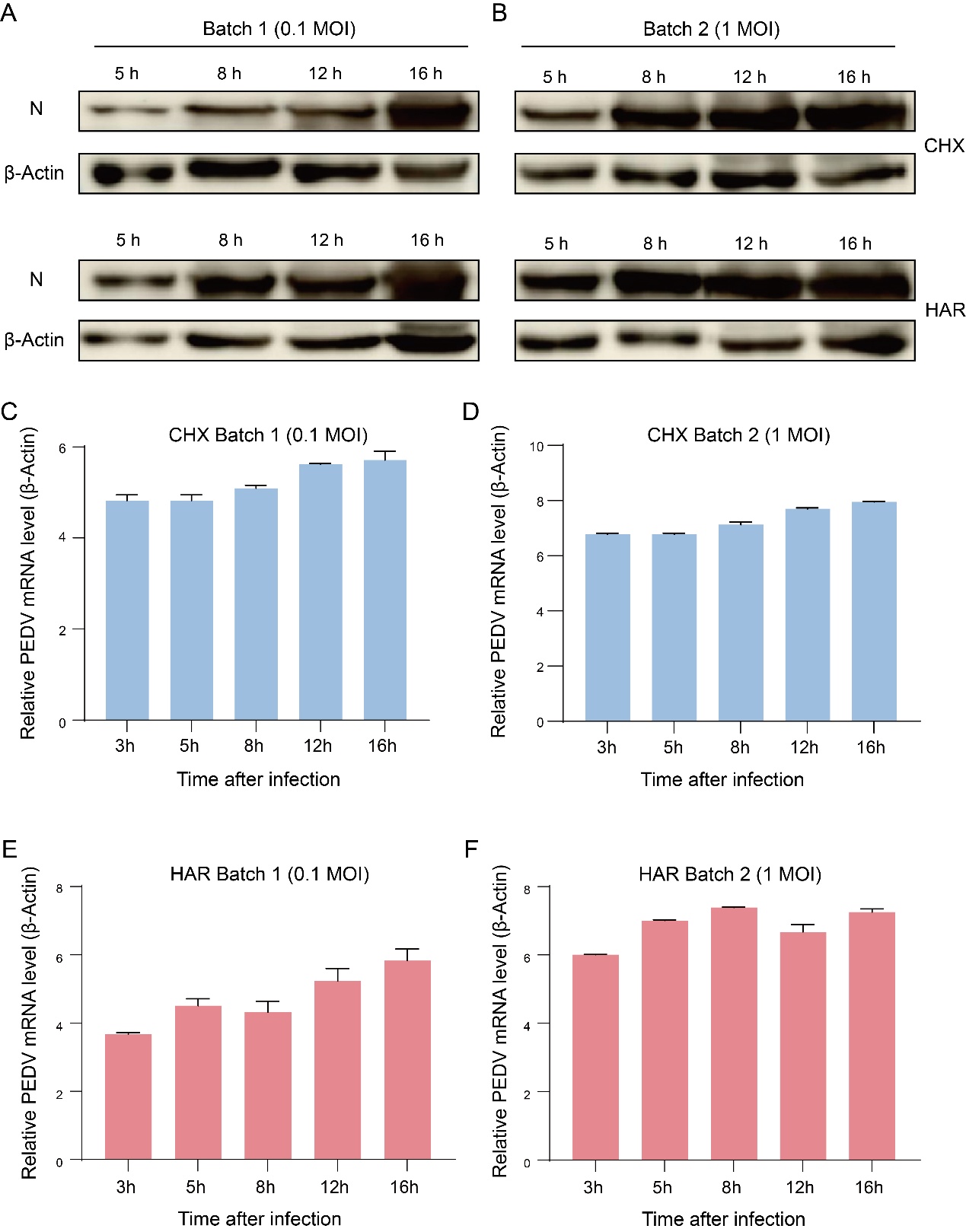

S1 Fig. Validation of PEDV infection by viral protein and RNA detection. (A, B) Western blot analysis of PEDV nucleocapsid (N) protein in infected Vero cells at 5, 8, 12, and 16 hours post-infection (hpi) under CHX and HAR treatment. Cells were infected at multiplicities of infection (MOIs) of 0.1 (batch 1) or 1 (batch 2). β-Actin served as the loading control. (C–F) RT-qPCR analysis of relative PEDV RNA levels at 3, 5, 8, 12, and 16 hpi under the indicated MOI and treatment conditions, normalized to β-actin. Error bars indicate variability among measurements.

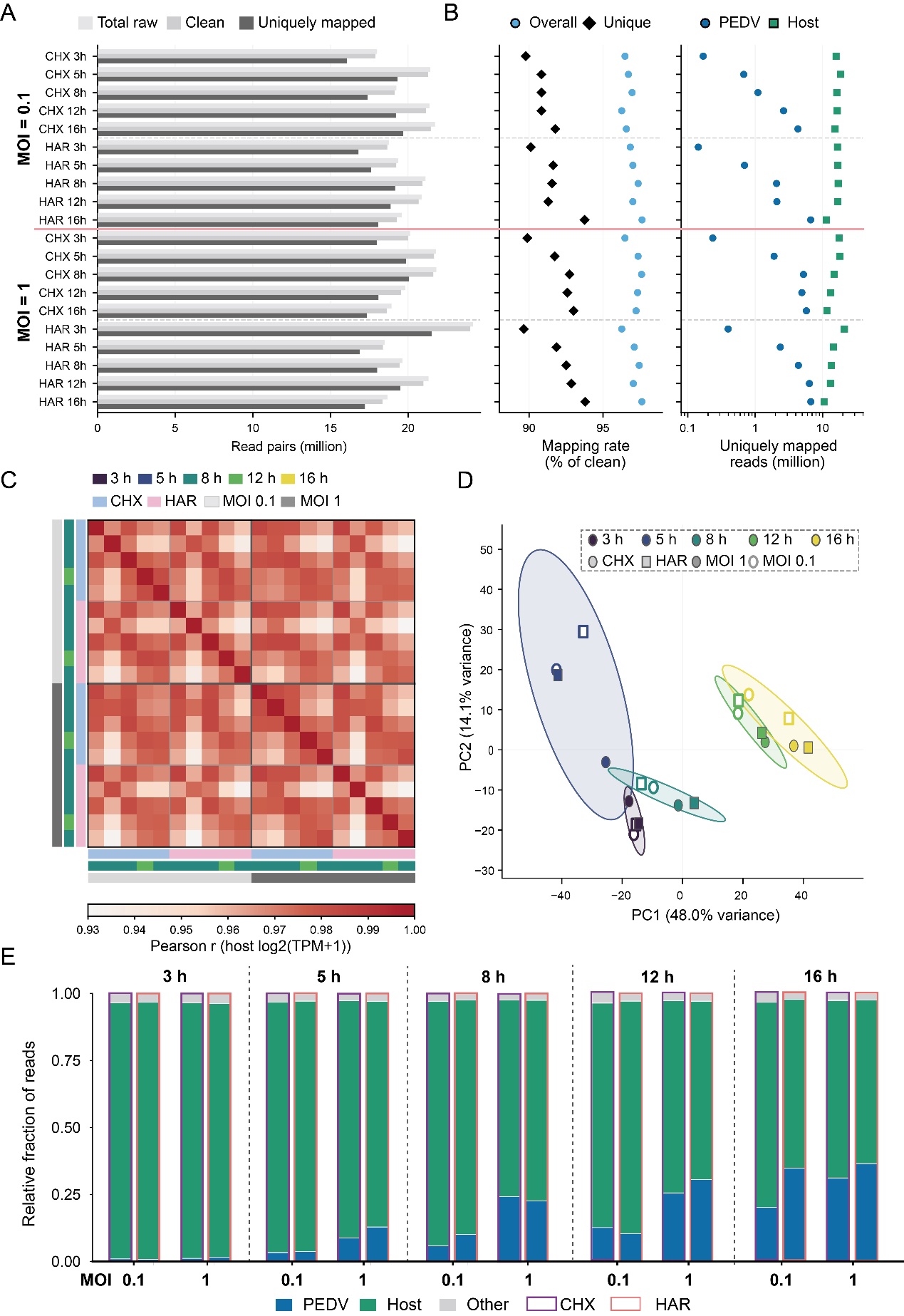

S2 Fig. RNA-seq quality assessment and sample relationships across PEDV infection conditions. (A) Numbers of raw, clean, and uniquely mapped sequencing read pairs for individual RNA-seq libraries. (B) Overall and unique mapping rates (left) and numbers of uniquely mapped PEDV and host reads (right). (C) Pairwise Pearson correlations of host gene expression profiles, calculated using log₂(TPM + 1). (D) Principal component analysis (PCA) of host gene expression profiles across infection time points, MOIs, and treatment conditions. PC1 and PC2 explain 48.0% and 14.1% of the variance, respectively. (E) Relative proportions of reads assigned to PEDV, host, and other sources in individual RNA-seq libraries.

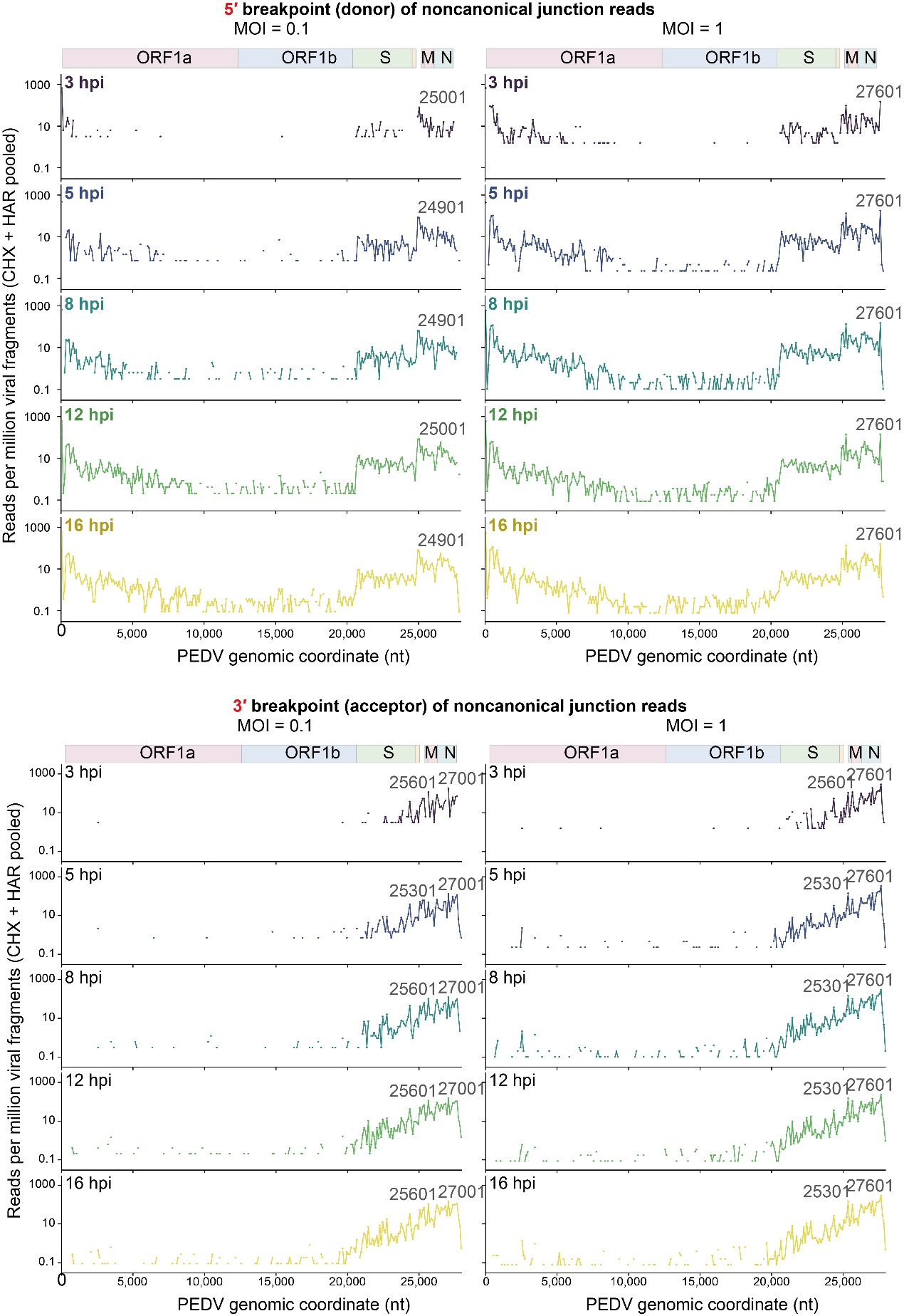

S3 Fig. Genome-wide breakpoint distributions of noncanonical PEDV RNA junctions across infection time points and MOIs. (A) Genomic distributions of 5′ breakpoints (donor sites) of noncanonical junction-supporting reads at 3, 5, 8, 12, and 16 hpi under MOI 0.1 (left) and MOI 1 (right). **(B)** Corresponding distributions of 3′ breakpoints (acceptor sites) under the same conditions. Read counts from CHX- and HAR-treated RNA-seq libraries were pooled and normalized per million viral fragments. Values are displayed on a logarithmic scale. Viral coding regions are annotated above each column, and labels indicate the genomic coordinates of prominent breakpoint peaks.

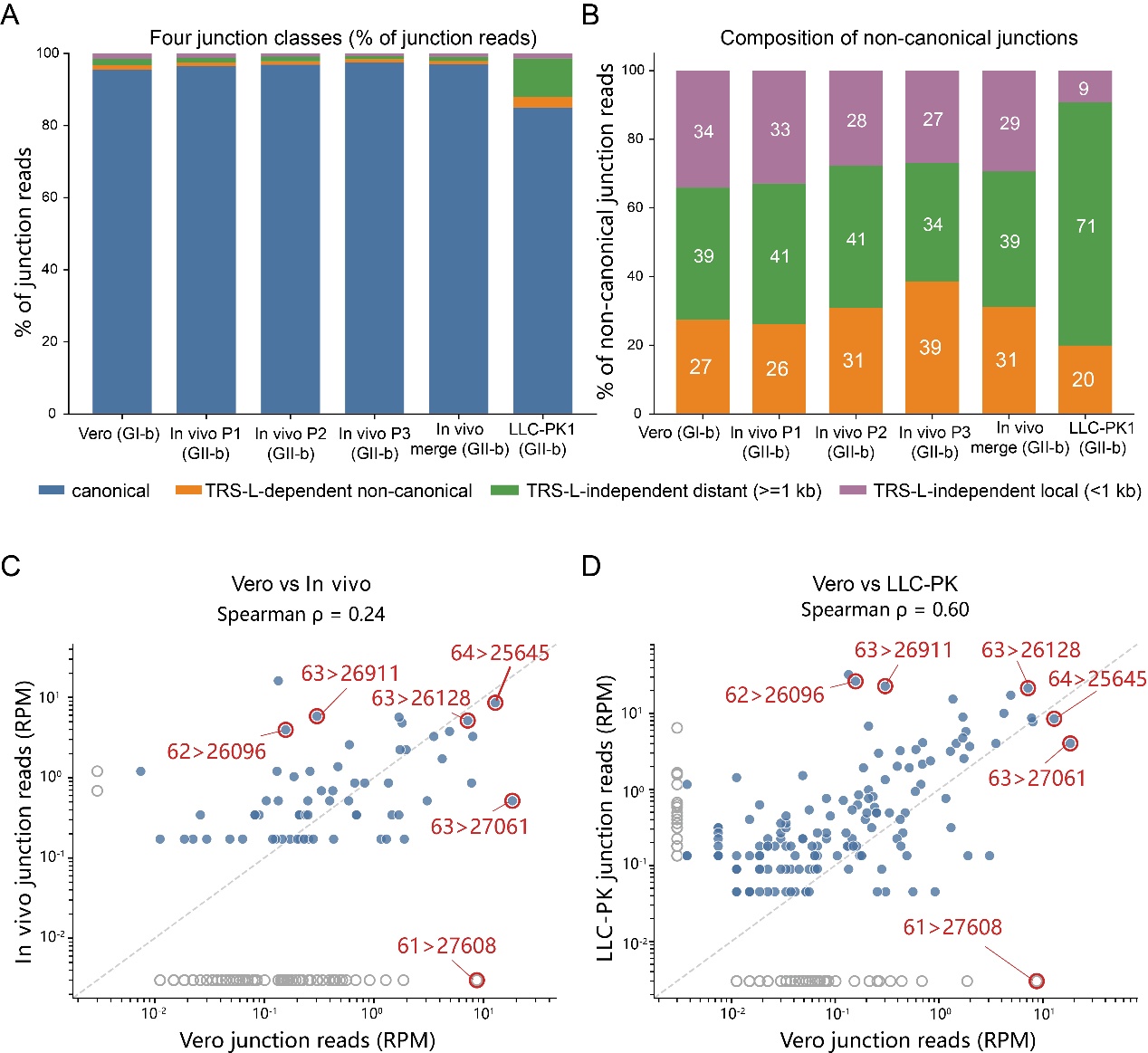

**S4 Fig. Comparison of PEDV RNA junction profiles across Vero, in vivo, and LLC-PK1 datasets.** (A) Relative proportions of four classes of RNA junctions in Vero GI-b, three individual in vivo GII-b samples, the merged in vivo dataset, and LLC-PK1 GII-b. Junctions were classified as canonical, TRS-L-dependent non-canonical, TRS-L-independent distant (≥1 kb), or TRS-L-independent local (<1 kb). (B) Relative composition of non-canonical junctions in the corresponding datasets, showing the contributions of TRS-L-dependent non-canonical, TRS-L-independent distant, and TRS-L-independent local junctions. (C, D) Scatter plots comparing TRS-L-dependent non-canonical junction-level read abundance (RPM) between Vero and in vivo datasets (C) and between Vero and LLC-PK1 datasets (D). Spearman correlation coefficients are indicated in each panel.

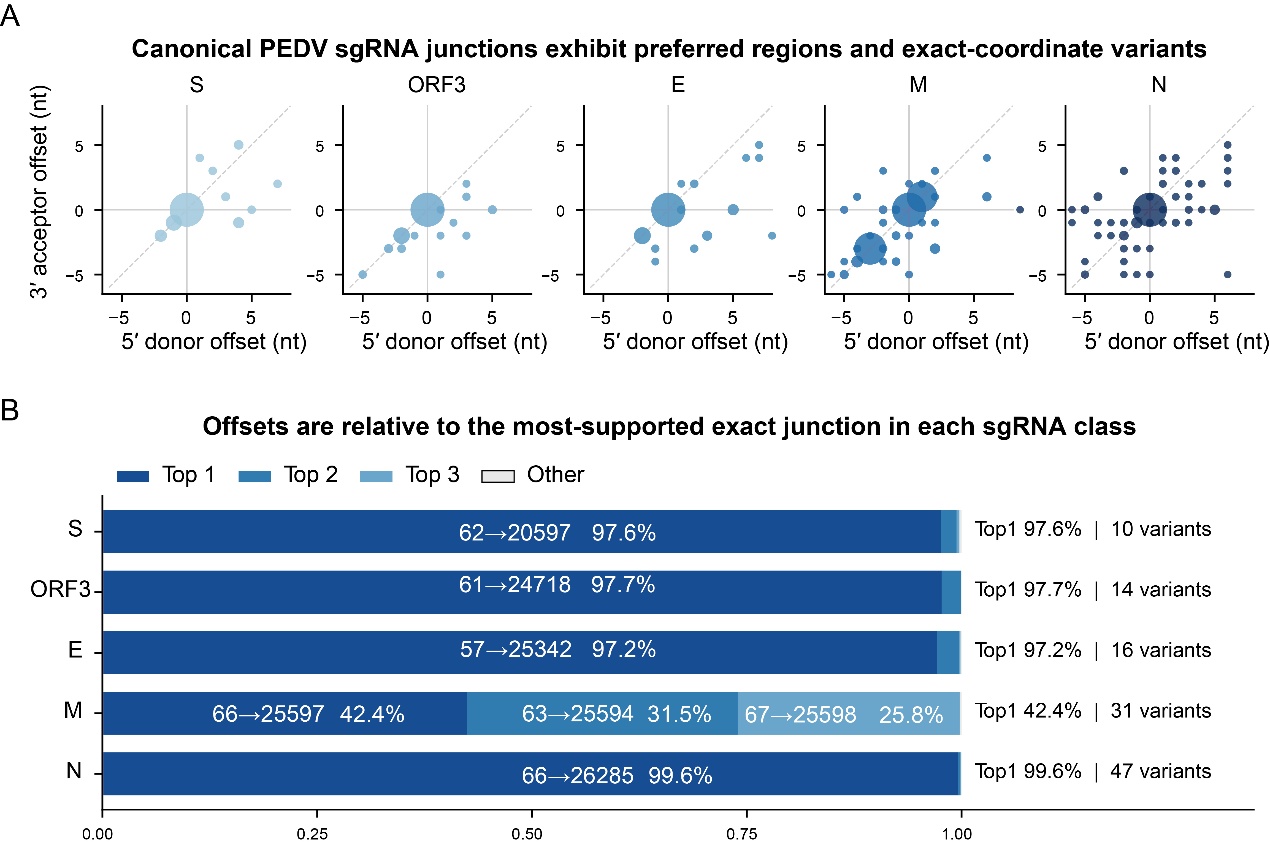

**S5 Fig. Canonical PEDV sgRNA junctions exhibit preferred breakpoint coordinates and local junction variants.** **(A)** Distributions of exact 5′ donor and 3′ acceptor coordinate offsets for the five canonical sgRNA classes (S, ORF3, E, M, and N). Offsets are calculated relative to the most abundant exact junction within each class. Bubble size represents the number of junction-supporting reads. **(B)** Relative contributions of the three most abundant exact junction variants and remaining variants within each canonical sgRNA class. Dominant junction coordinates, their read fractions, and the total numbers of exact variants are indicated.

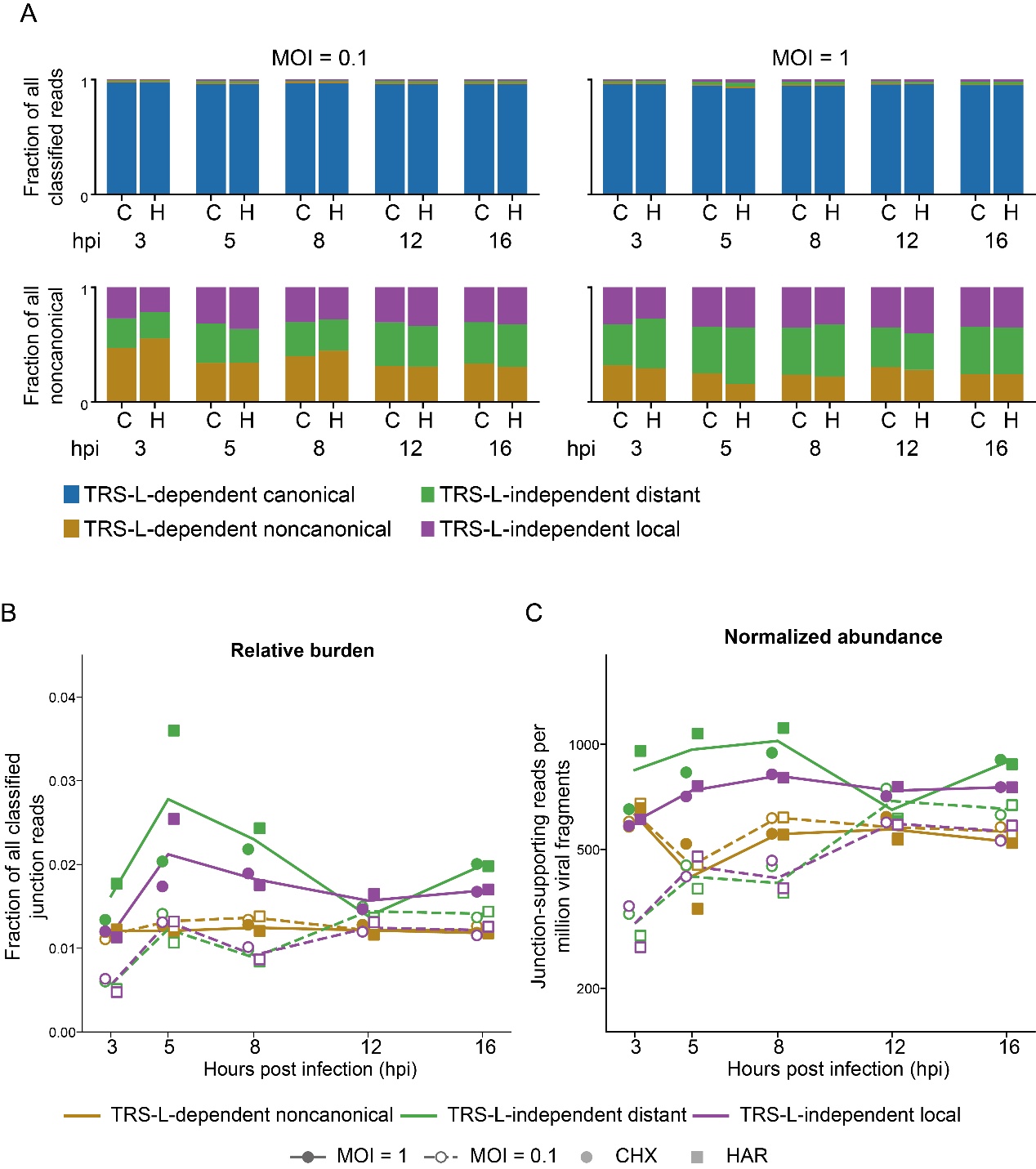

**S6 Fig. Temporal changes in canonical and noncanonical PEDV RNA junction classes across infection conditions.** **(A)** Relative proportions of canonical and noncanonical junction classes among all classified junction-supporting reads (upper panels), and the composition of noncanonical junction reads (lower panels), at 3, 5, 8, 12, and 16 hours post-infection (hpi). Data are shown separately for multiplicities of infection (MOIs) of 0.1 and 1 and for cycloheximide (C)- and harringtonine (H)-treated libraries. **(B)** Temporal changes in the relative burden of noncanonical junction classes, expressed as fractions of all classified junction-supporting reads (left), and their abundance normalized per million viral fragments (right). Colors indicate junction classes; line styles indicate MOI, and symbols indicate treatment.

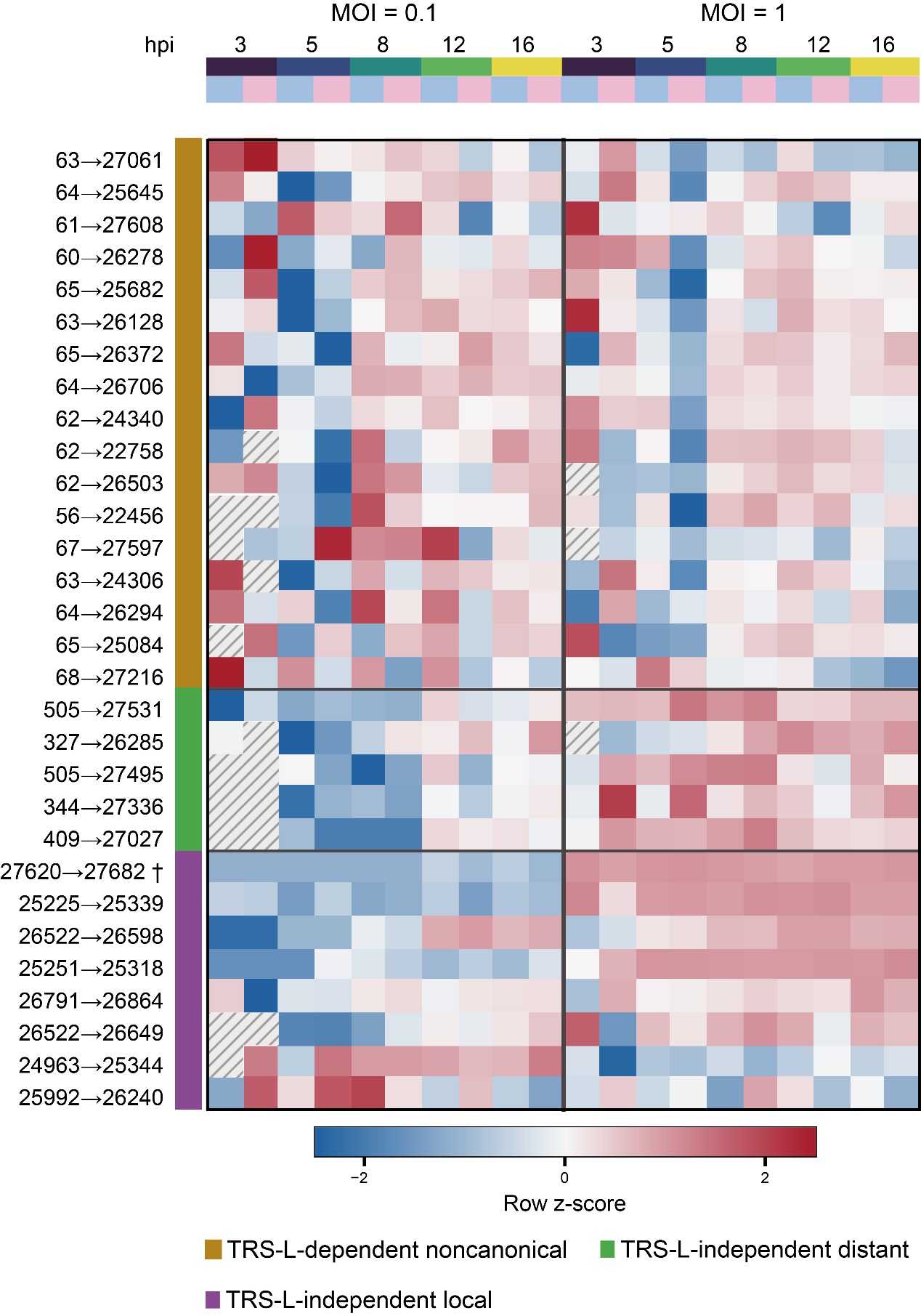

**S7 Fig. Temporal profiles of representative noncanonical PEDV RNA junctions across infection conditions.**

Heatmap showing the relative abundance of selected noncanonical junctions at 3, 5, 8, 12, and 16 hpi under MOIs of 0.1 and 1 in CHX- and HAR-treated RNA-seq libraries. Rows represent individual junctions identified by their exact 5′ donor and 3′ acceptor coordinates, grouped as TRS-L-dependent noncanonical, TRS-L-independent distant, or TRS-L-independent local junctions. Values are row-wise z-scores. Hatched cells indicate unavailable values.

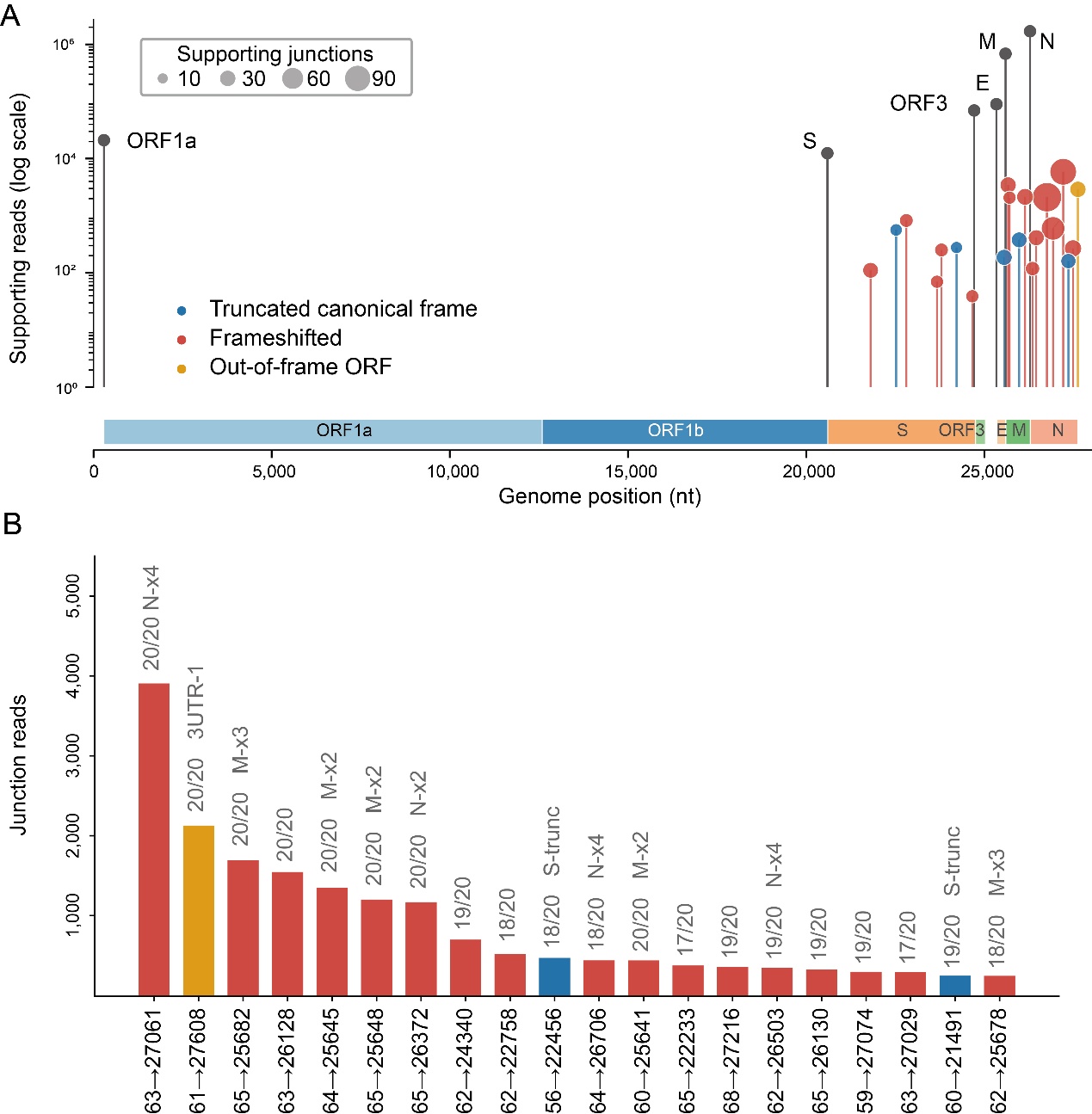

**S8 Fig. Abundant noncanonical PEDV leader–body junctions exhibit distinct predicted coding consequences and sequence features.** **(A)** Genomic distribution and read abundance of canonical and selected noncanonical leader–body junctions. Noncanonical junctions are colored according to their predicted coding consequences, including truncated canonical coding sequences, junction-associated frameshifted ORFs, and out-of-frame ORFs. Bubble sizes indicate the numbers of supporting junctions. **(B)** The 20 most abundant TRS-L-dependent noncanonical junctions, ranked by junction-supporting read counts. Exact breakpoint coordinates, sample detection frequencies, and selected predicted coding products are indicated.

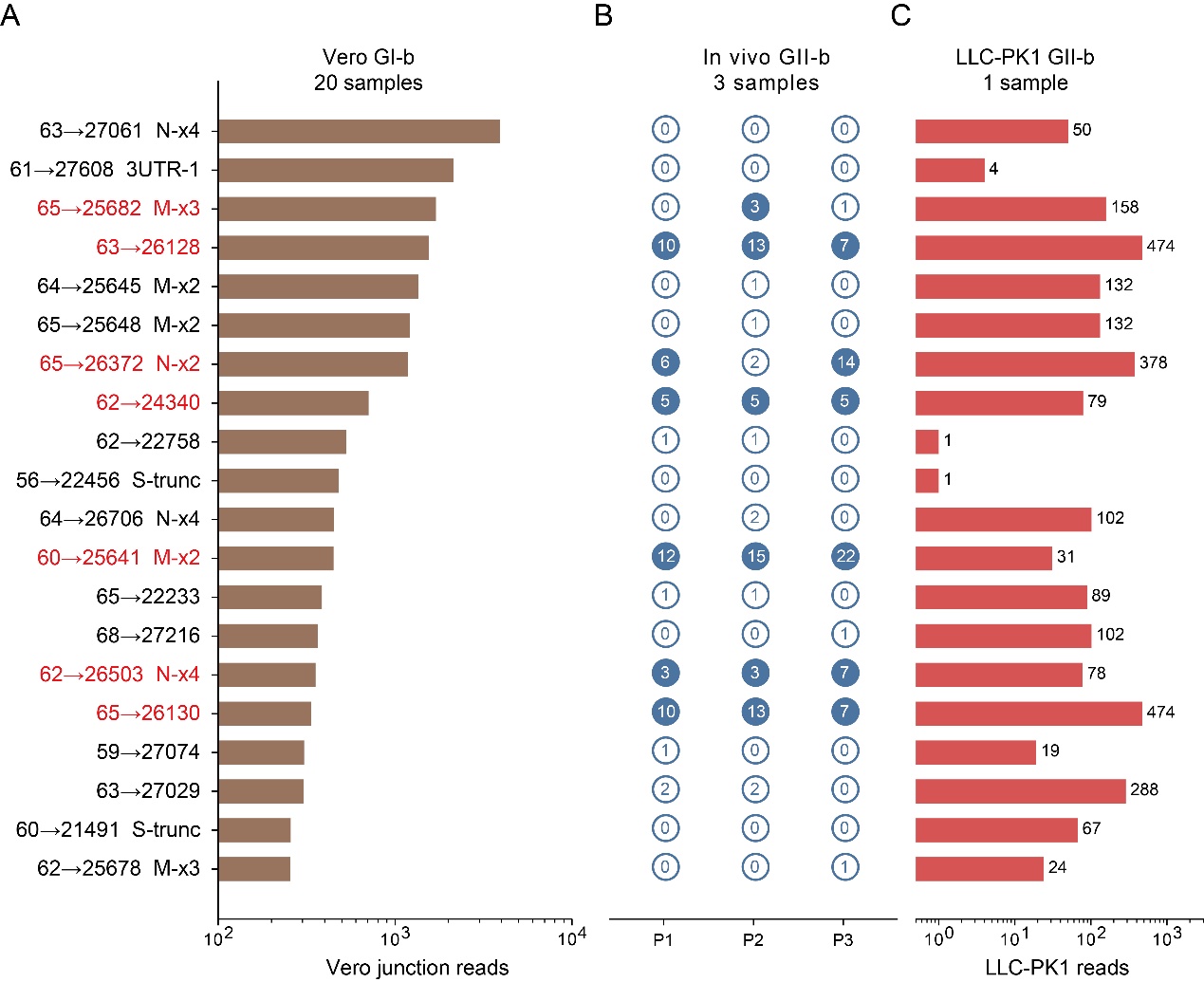

**S9 Fig. Detection and abundance of selected PEDV RNA TRS-L-dependent non-canonical junctions across Vero, in vivo, and LLC-PK1 datasets.** (A) Junction-read abundance of the 20 selected PEDV RNA junctions in the Vero GI-b dataset, comprising 20 samples. Junctions are ranked according to their read abundance, with corresponding genomic coordinates and junction annotations indicated on the left. (B) Detection of the same junctions in three publicly available in vivo GII-b samples (P1–P3). The numbers inside the circles indicate junction-read counts in each sample; filled and open circles distinguish detected and undetected junctions, respectively. (C) Junction-read abundance of the corresponding junctions in one publicly available LLC-PK1 GII-b dataset. Numbers adjacent to the bars indicate read counts.

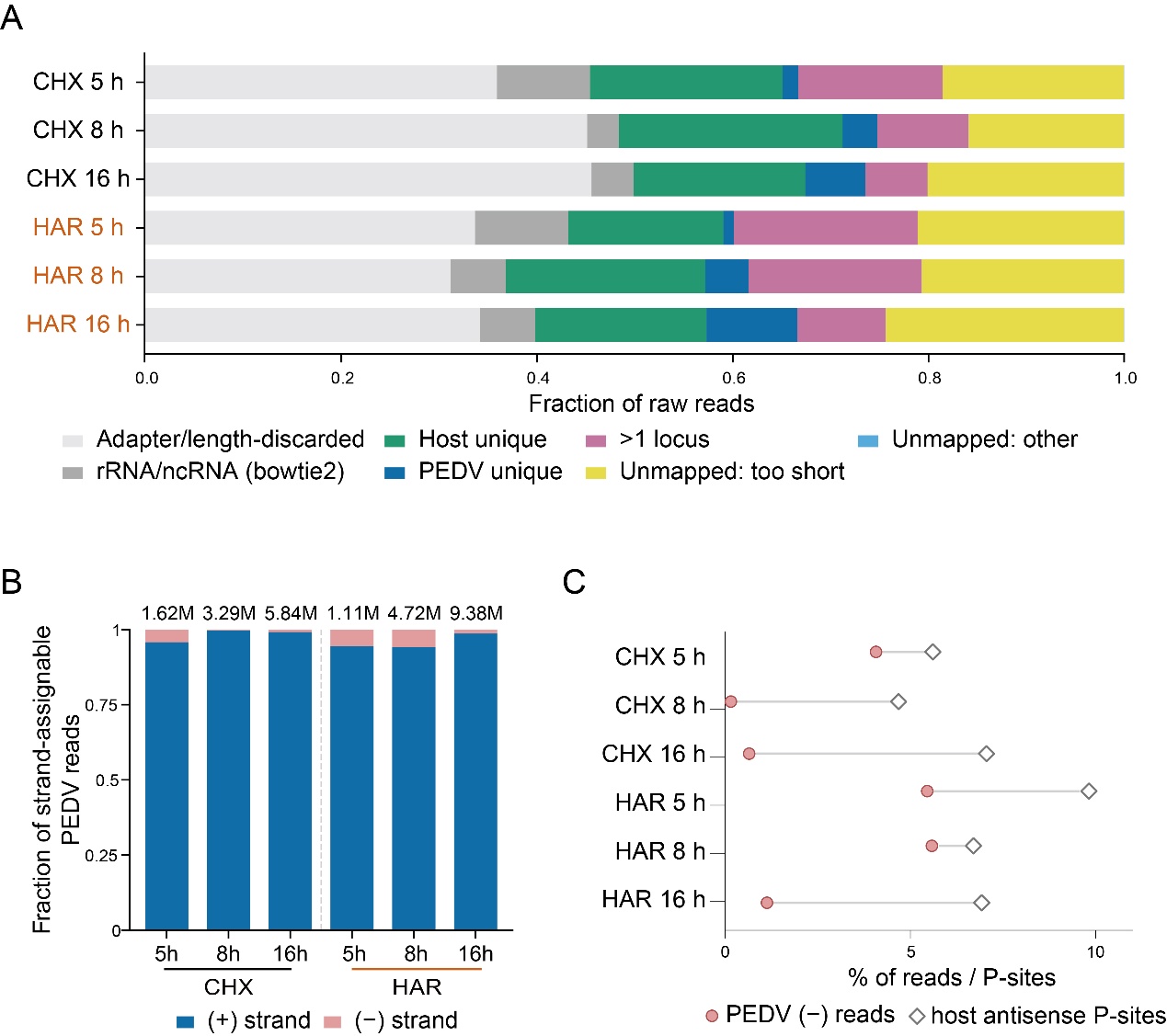

**S10 Fig. Read processing, strand distribution, and antisense signals in PEDV ribosome profiling libraries.** **(A)** Fractions of raw reads assigned to successive filtering and mapping categories in CHX- and HAR-treated Ribo-seq libraries at 5, 8, and 16 hours post-infection (hpi). **(B)** Strand distribution of PEDV-aligned reads, showing the proportions of positive- and negative-strand reads. Numbers above the bars indicate total strand-assignable viral reads. **(C)** Percentages of PEDV negative-strand reads and host antisense P-sites across the six Ribo-seq libraries. **(D)** Median aligned footprint lengths of host- and PEDV-derived reads in CHX- and HAR-treated libraries at 5, 8, and 16 hpi. Connected symbols indicate matched infection time points; numbers indicate the PEDV–host differences in median length. **(E)** Fractions of aligned footprints measuring 29–31 nucleotides in the corresponding host and PEDV libraries. Labels indicate the differences in fractions between PEDV and host.

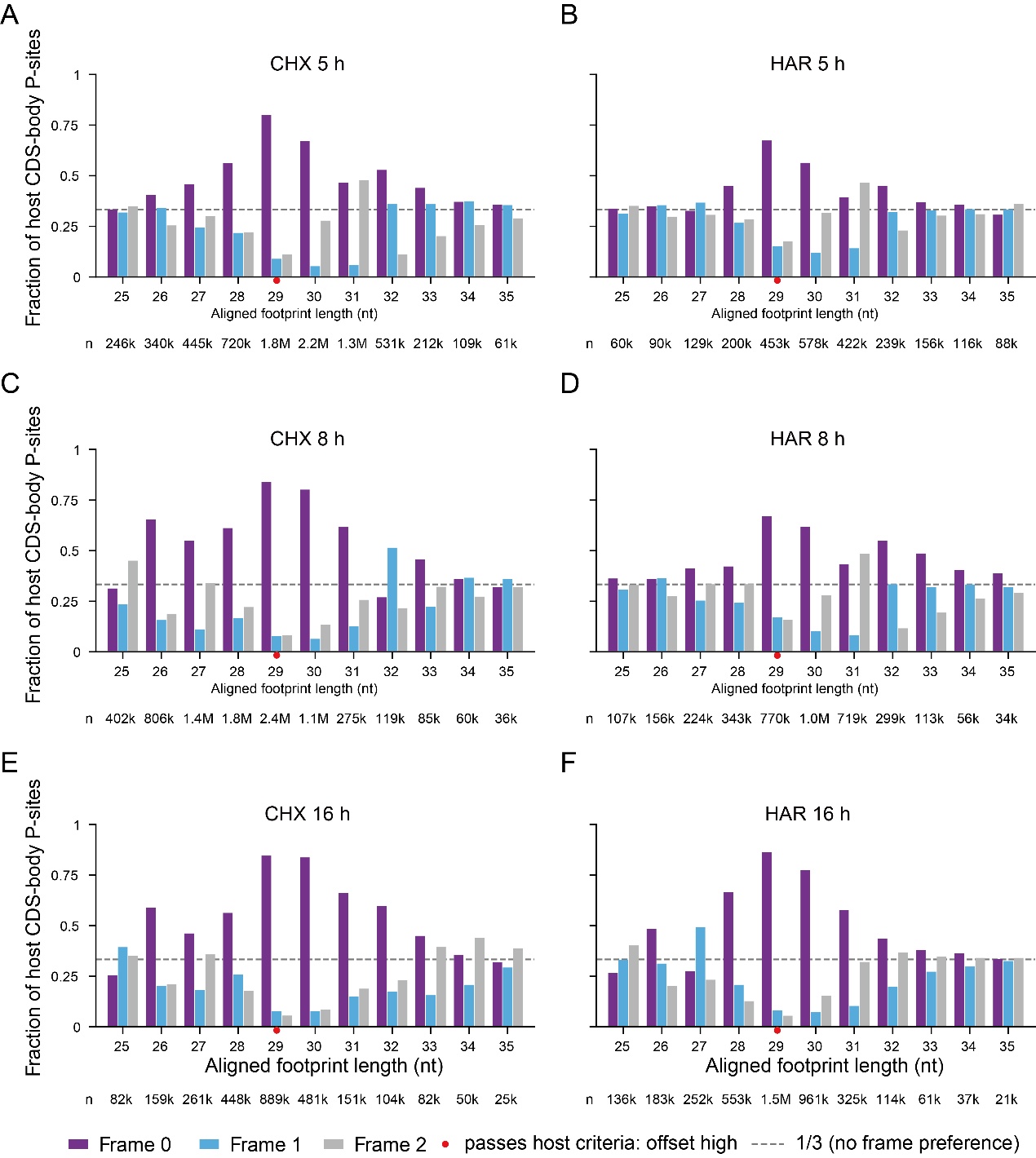

**S11 Fig. Footprint length-dependent reading-frame periodicity supports the selection of 29-nt RPFs for alternative ORF analysis.** **(A–F)** Reading-frame distributions of host coding sequence (CDS)-body P-sites across aligned footprint lengths of 25–35 nt in CHX- and HAR-treated libraries at 5, 8, and 16 hpi. Purple, blue, and gray bars represent frame 0, frame +1, and frame +2, respectively. The dashed horizontal line indicates the expected fraction of one-third in the absence of reading-frame preference. Read counts are shown below each footprint length. Red dots mark 29-nt footprints that pass the host-based P-site offset criteria.

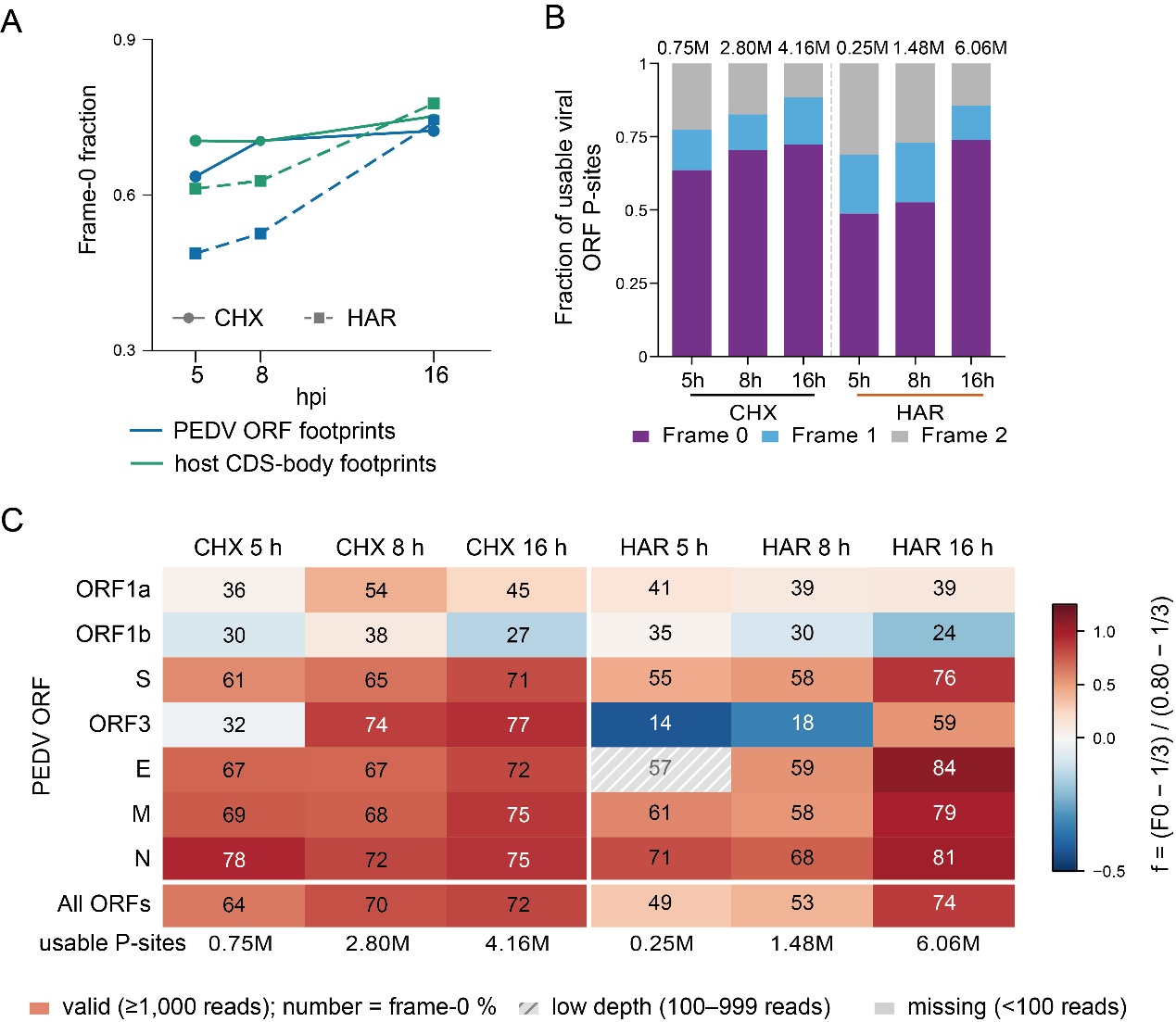

**S12 Fig. Temporal and gene-specific variation in PEDV reading-frame occupancy.** **(A)** Fraction of P-sites assigned to the annotated frame 0 in PEDV ORFs and host CDS bodies across CHX- and HAR-treated libraries at 5, 8, and 16 hpi. **(B)** Reading-frame composition of usable PEDV ORF P-sites in the six libraries. Numbers above the bars indicate total usable P-sites. **(C)** Heatmap of frame-0 occupancy across individual PEDV ORFs and infection conditions. Cell labels indicate the percentage of P-sites assigned to frame 0. Colors represent deviation from the expected one-third frame occupancy, scaled relative to a reference fraction of 0.80. Hatched and gray cells indicate low-depth (100–999 reads) and insufficient-depth (<100 reads) measurements, respectively.

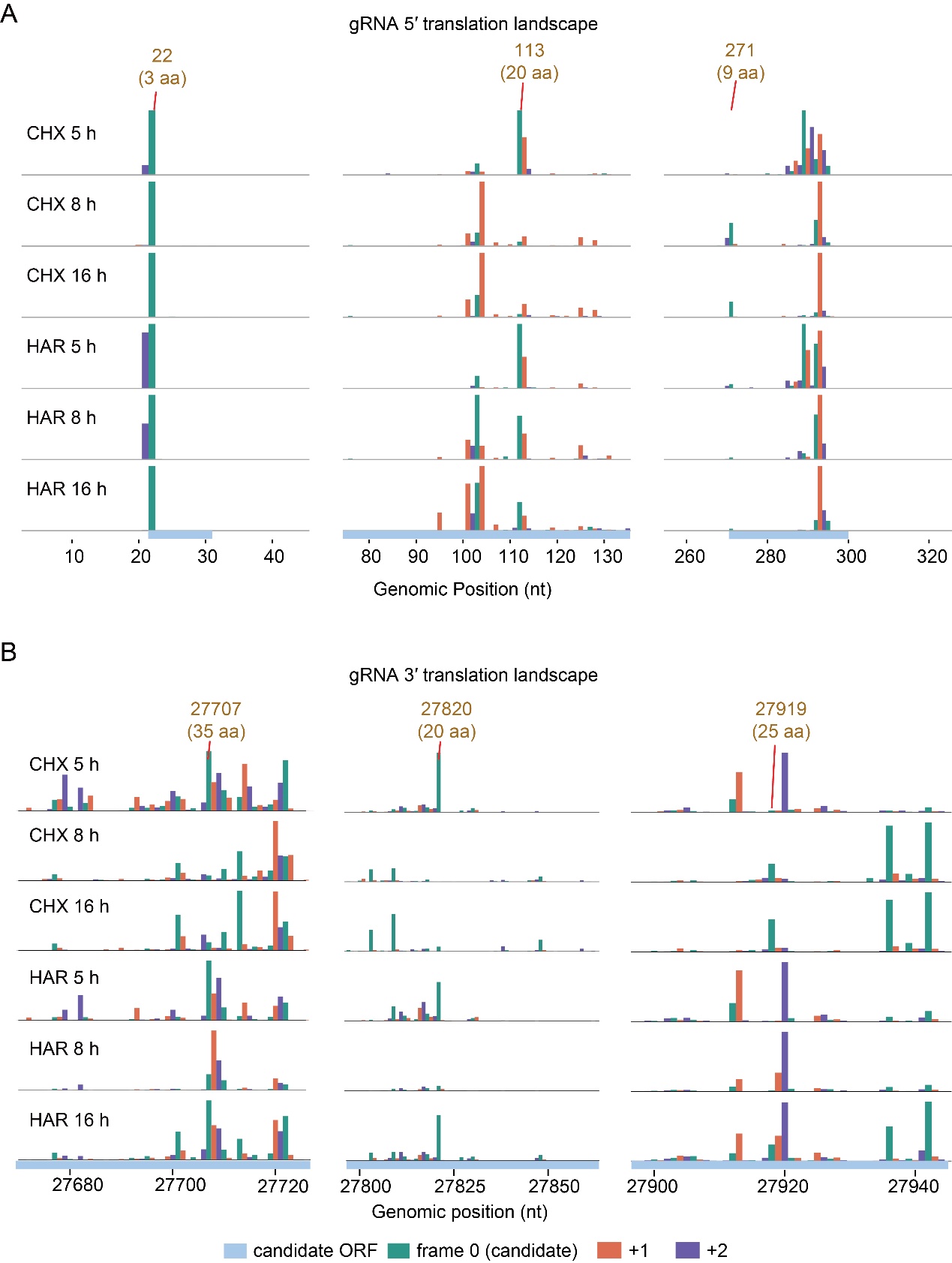

**S13 Fig. Frame-resolved ribosome occupancy reveals localized translation-associated signals in the PEDV 5′ and 3′ untranslated regions.** **(A)** P-site distributions around three major hotspots in the genomic 5′ region (positions 22, 113, and 271 nt). **(B)** Corresponding P-site distributions around three hotspots in the genomic 3′ region (positions 27,707, 27,820, and 27,919 nt).

Profiles are shown for cycloheximide (CHX)- and harringtonine (HAR)-treated libraries at 5, 8, and 16 hours post-infection (hpi). Colors indicate P-site reading frames relative to the candidate ORFs: frame 0 (green), +1 (orange), and +2 (purple). Light-blue bars indicate candidate ORF intervals. Numbers above the hotspots indicate genomic positions and predicted peptide lengths.

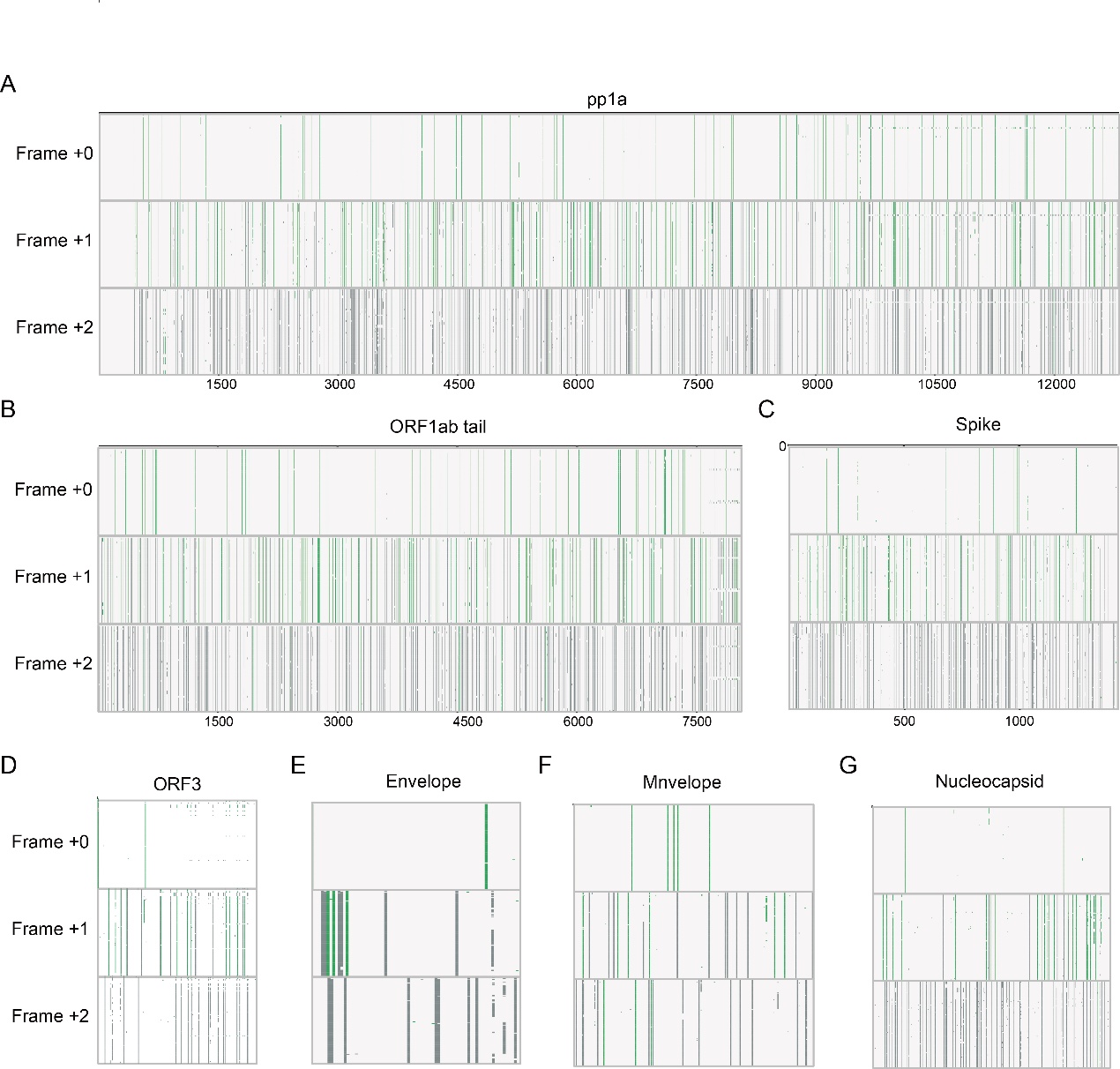

**S14 Fig. Genome-wide distribution of overlapping reading-frame sequence features across annotated PEDV genes.** **(A–G)** Distribution of sequence features in the annotated reading frame (0) and the alternative +1 and +2 reading frames across PEDV coding regions: ORF1a, ORF1b, S, ORF3, E, M, and N. Vertical marks indicate the genomic positions of the corresponding sequence features in each reading frame.

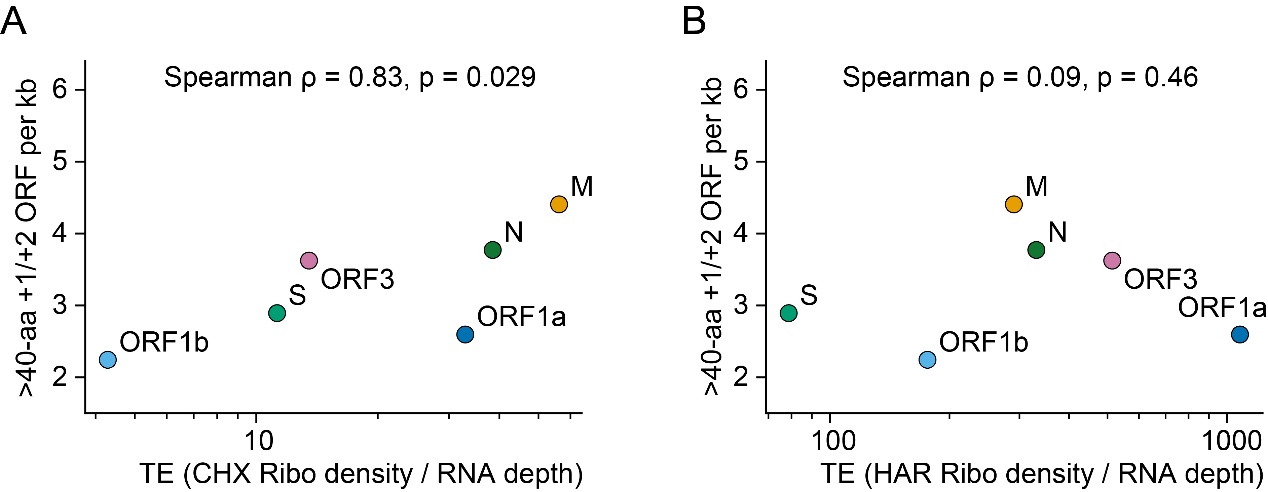

**S15 Fig. Density of long overlapping ORFs is associated with CHX-based translation efficiency across PEDV coding regions.** **(A)** Relationship between the density of overlapping +1/+2 ORFs longer than 40 amino acids per kilobase and translation efficiency (TE), calculated as CHX Ribo-seq density divided by RNA-seq depth. **(B)** Corresponding relationship using HAR Ribo-seq density divided by RNA-seq depth. Each point represents an annotated PEDV coding region. Spearman correlation coefficients and P values are shown in the panels.

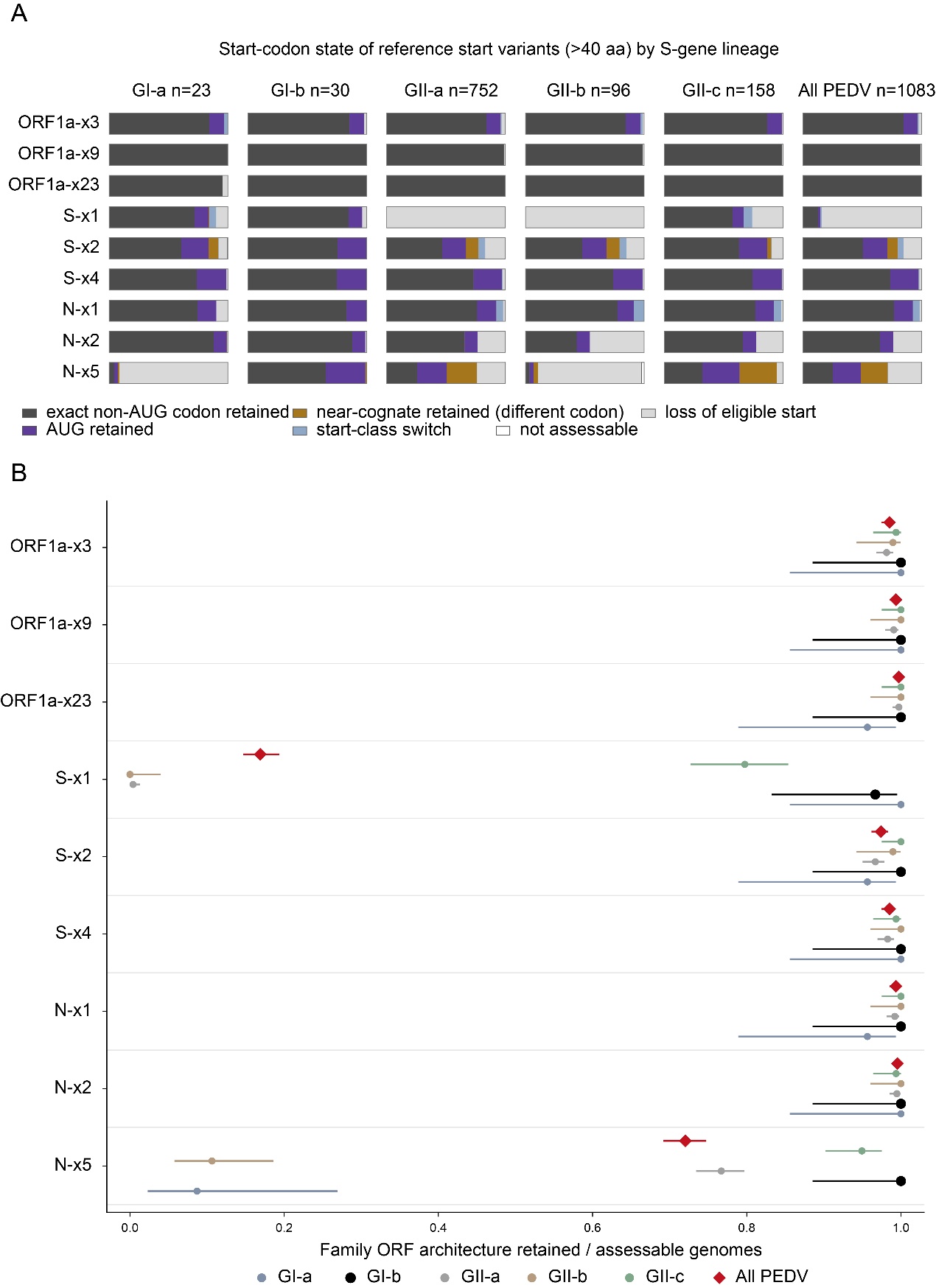

**S16 Fig. Evolutionary conservation of candidate alternative open reading frames across PEDV lineages.** (A) Conservation of predicted initiation codons for representative alternative ORFs longer than 40 amino acids across PEDV lineages. Complete PEDV genome sequences were retrieved from the NCBI database, deduplicated, and classified into five S-gene lineages (GI-a, GI-b, GII-a, GII-b, and GII-c). Initiation codons were classified as exact non-AUG codon retained, AUG retained, alternative near-cognate codon retained, start-codon class switched, loss of eligible initiation codon, or not assessable. The number of genomes in each lineage is indicated above the corresponding columns. (B) Proportion of assessable genomes retaining the reference-defined ORF architecture for each candidate alternative ORF across the five PEDV lineages and the complete dataset (n = 1,083). Points indicate the proportion of genomes retaining the predicted ORF architecture, with horizontal intervals showing the corresponding uncertainty estimates. Colors distinguish viral lineages, and red diamonds represent the combined PEDV dataset.

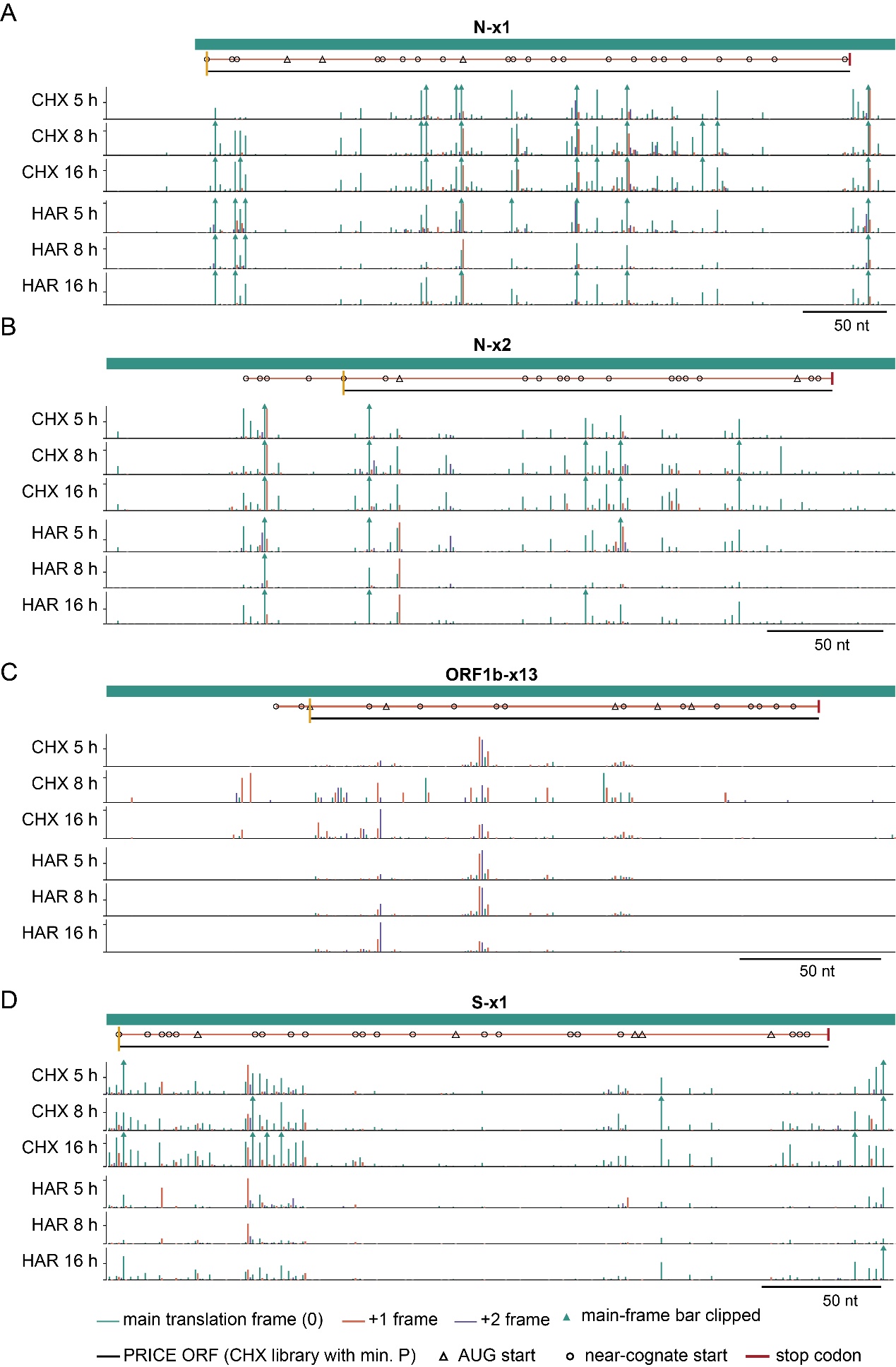

**S17 Fig. Frame-resolved ribosome profiling supports selected candidate overlapping ORFs in PEDV.** **(A–D)** Local P-site distributions surrounding four representative candidate overlapping ORFs: N-x1, N-x2, ORF1b-x13, and S-x1. CHX- and HAR-treated libraries at 5, 8, and 16 hpi are shown separately. P-sites are colored according to their reading frames relative to the annotated parental ORF (frame 0, green; +1, orange; +2, purple). Candidate ORF structures and initiation codons are indicated above each profile. Black lines represent ORFs identified by PRICE using CHX data, with the selected initiation site indicated by a yellow vertical mark. Triangles and open circles denote AUG and near-cognate initiation codons, respectively; red terminal marks indicate stop codons. Green triangles indicate clipped main-frame P-site peaks. Scale bars represent 50 nt.

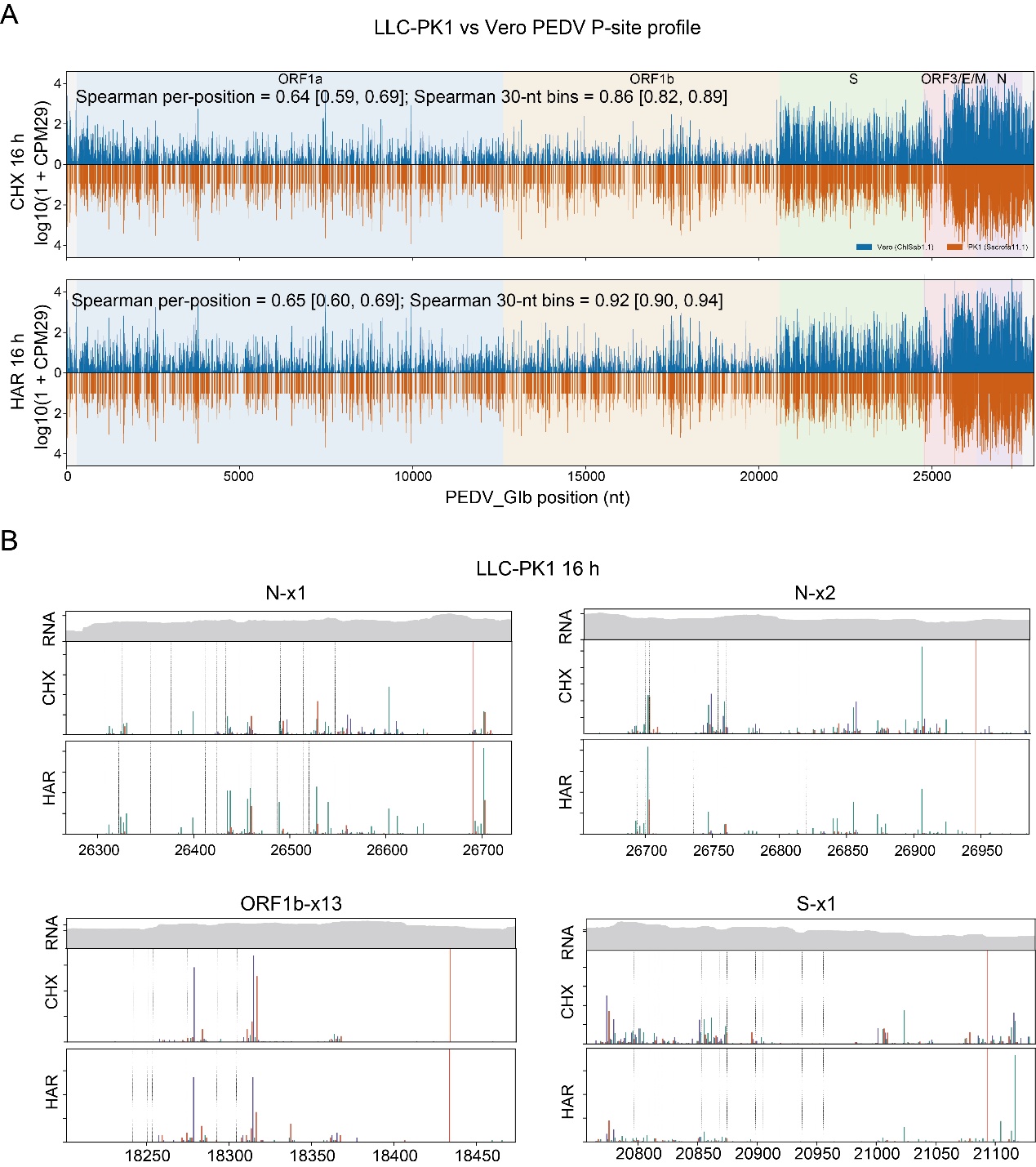

**S18 Fig. Comparison of genome-wide PEDV P-site profiles between Vero and LLC-PK1 cells at 16 h post-infection.** (A) Genome-wide comparison of normalized P-site densities between Vero and LLC-PK1 datasets under cycloheximide (CHX) and harringtonine (HAR) treatment at 16 h post-infection (hpi). P-site densities are expressed as log10 (1 + CPM29), with Vero and LLC-PK1 profiles shown in blue and orange, respectively. Spearman correlation coefficients are provided for individual nucleotide positions and 30-nt binned profiles, together with their corresponding 95% confidence intervals. (B) Local RNA coverage and P-site distributions under CHX and HAR conditions across four representative PEDV candidate alternative ORF regions: N-x1, N-x2, ORF1b-x13, and S-x1. Gray profiles represent RNA coverage, while the lower tracks display the corresponding CHX- and HAR-associated P-site distributions.

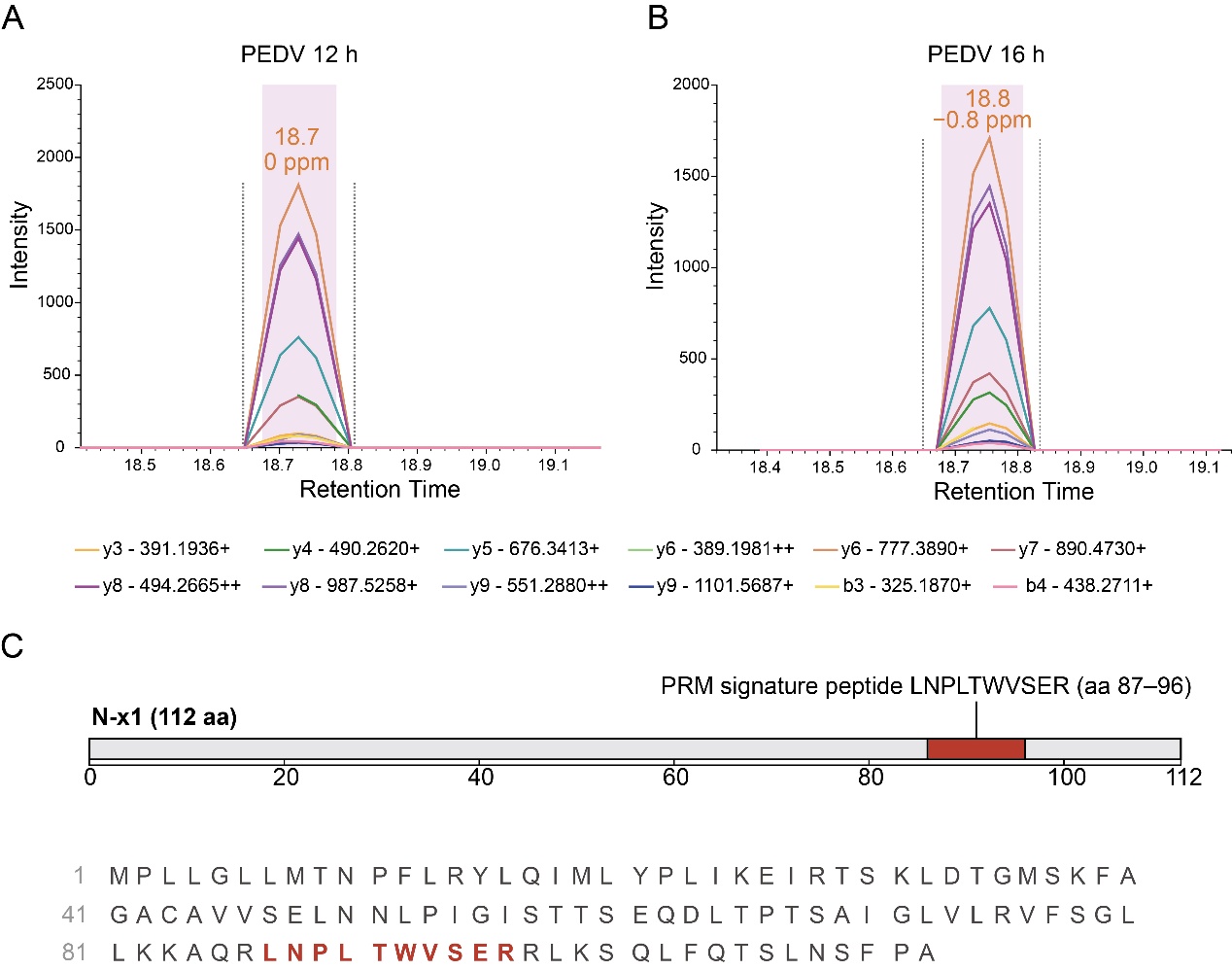

**S19 Fig. Parallel reaction monitoring (PRM) analysis of the candidate N-x1-encoded signature peptide in PEDV-infected cells.** (A, B) Representative PRM chromatograms of the signature peptide LNPLTWVSER (amino acids 87–96) detected in PEDV-infected samples at 12 h and 16 h, respectively. Multiple fragment-ion transitions, including y- and b-series ions, are displayed to assess peptide-specific detection and retention-time consistency. (C) Schematic representation of the predicted 112-amino-acid N-x1 protein, indicating the position of the PRM signature peptide LNPLTWVSER (amino acids 87–96), highlighted in red.
